# Discovery of a Novel Polymer for Xeno-free, Long-term Culture of Human Pluripotent Stem Cell Expansion

**DOI:** 10.1101/2020.09.16.298810

**Authors:** Jordan Thorpe, Aishah Nasir, Laurence Burroughs, Joris Meurs, Sara Pijuan-Galito, Derek J. Irvine, Morgan R. Alexander, Chris Denning

## Abstract

Human pluripotent stem cells (hPSCs) can be expanded and differentiated *in vitro* into almost any adult tissue cell type, and thus have great potential as a source for cell therapies with biomedical application. In this study, a fully-defined polymer synthetic substrate is identified for hPSC culture in completely defined, xeno-free conditions. This system can overcome the cost, scalability and reproducibility limitations of current hPSC culture strategies, and facilitate large-scale production. A high-throughput, multi-generational polymer microarray platform approach was used to test over 600 unique polymers and rapidly assess hPSC-polymer interactions in combination with the fully defined xeno-free medium, Essential 8^TM^ (E8). This study identifies as novel nanoscale phase separated blend of poly(tricyclodecane-dimethanol diacrylate) and poly(butyl acrylate) (2:1 v/v), which supports long-term expansion of hPSCs and can be readily coated onto standard cultureware. Analysis of cell-polymer interface interactions through mass spectrometry and integrin blocking studies provides novel mechanistic insight into the role of the E8 proteins in promoting integrin-mediated hPSC attachment and maintaining hPSC signaling, including ability to undergo multi-lineage differentiation. This study therefore identifies a novel substrate for long-term serial passaging of hPSCs in serum-free, commercial chemically-defined E8, which provides a promising and economic hPSC expansion platform for clinical-scale application.

## Main section

To improve application of adherent hPSC culture, there has been a shift from the use of animal-derived feeder layers and poorly defined media to systems using surfaces and culture media that are fully-defined.^[1–7]^ High-throughput material screening platforms have facilitated discovery of synthetic culture surfaces for commercialization, including the fully defined peptide-containing surface, Synthemax^TM^ (Corning ®), which offer alternative growth substrates to widely-used but poorly defined mouse sarcoma preparation, Matrigel^TM^.^[8]^ Peptide-containing surfaces still require incorporation of biologically active motifs (derived from extracellular matrix proteins) to maintain long-term hPSC culture (at least 8 passages), which significantly increase costs and limit use for large-scale production (estimated at $10,000-$15,000 for 1 billion hPSCs required for a single patient intervention).^[9]^

Use of polymers as culture substrates, is a safe and cost-effective solution. Previous analysis of hPSC-polymer interactions, assessed using high-throughput polymer screening strategies, demonstrated the influence of extracellular matrix components (eg. fibronectin) and/or xenogeneic components from culture medium (eg. bovine serum albumin containing commercial defined mTESR1 and StemPro) for maintaining hPSC culture on synthetic surfaces.^[10, 11]^ In the following study, we report the discovery of novel materials capable of supporting hPSCs in a culture system simplified by using the xeno-free, commercial, chemically-defined E8^TM^ medium. Subsequent characterization of cell interactions with the scaled up synthetic polymer revealed the underlying mechanisms governing cellular response without the contribution of xenogeneic components.

Using a multigenerational polymer microarray screening approach, the first generation array consisted of 284 chemically diverse monomers (photo-curable and readily commercially available) pin-printed and UV polymerized onto poly(2-hydroxyethyl methacrylate) (poly(HEMA)) coated slides as spots in triplicate (**Figure 1**a, monomer structures presented in Figure S1 with monomer names in Table S1).^[12, 13]^ ReBl-PAT cells, a human induced pluripotent stem cell (hiPSC) line generated in-house,^[14]^ were seeded onto arrays in E8 for 24 h before quantification of cell attachment and response using automated fluorescence microscopy. Materials were ranked by total number of nuclei for cell attachment gauged by DAPI staining (Figure S2a, see rank order with polymer names Table S2), or by number of OCT4+ cells as a marker of pluripotency (Figure S2b, see rank order with polymer names Table S2), and the two parameters were plotted against each other (Figure 1b, c).

**Figure 1.**
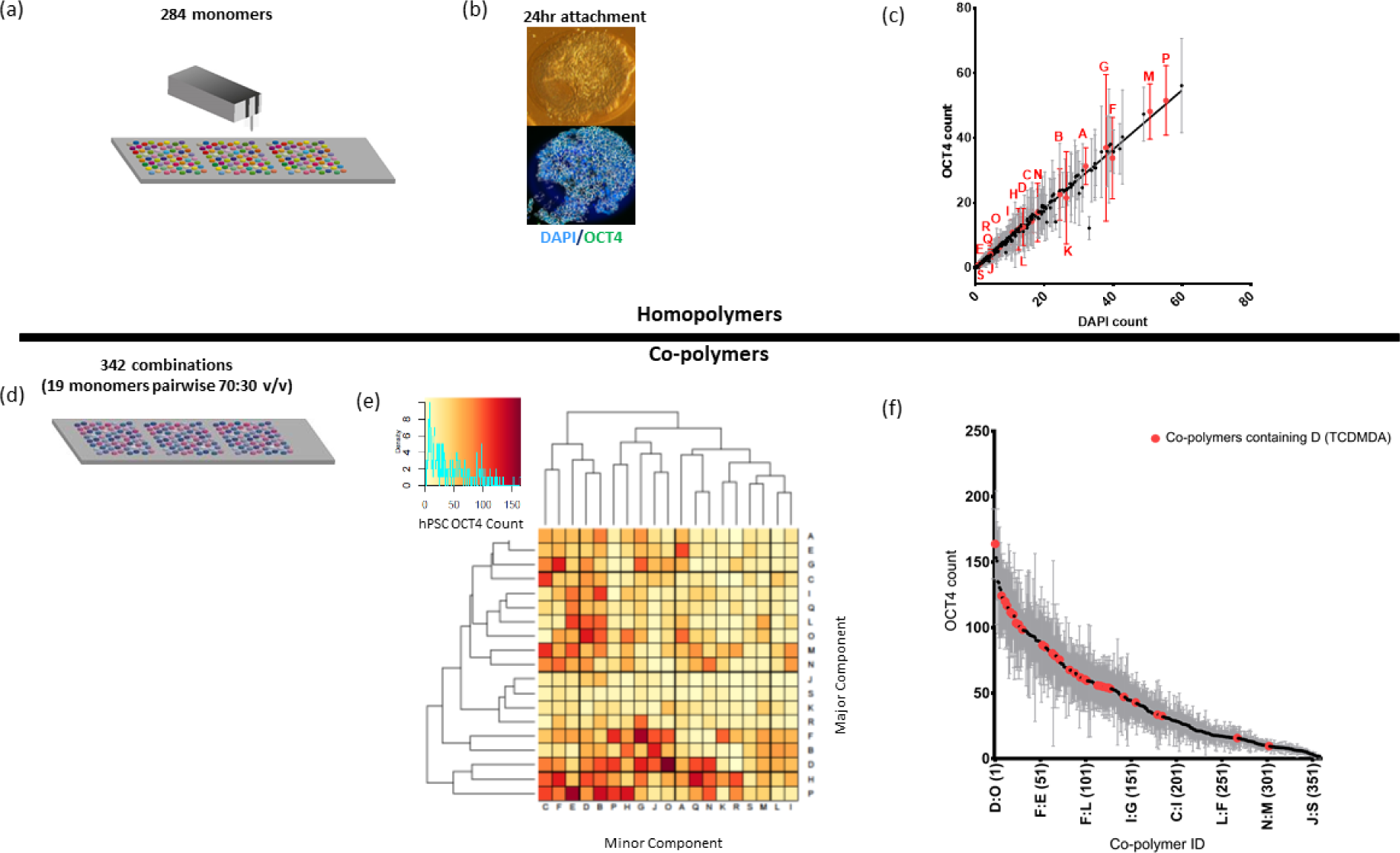
Multi-generation microarray screen of polymeric substrates. (a) A first-generation array of 284 chemically diverse monomers were screened for hPSC attachment with ReBl-PAT hiPSCs in E8 medium for 24 h. (b) Arrays were then fixed and stained for pluripotent marker OCT4, imaged using Imstar automated fluorescence microscopy and OCT4+ nuclei and total nuclei counts assessed with Cell-Profiler. Representative image shows a polymer spot (n) supporting high hPSC attachment. (c) Attachment on materials are ranked by OCT4+ nuclei count plotted against total cell number (DAPI). Nineteen materials selected for second-generation co-polymer screening (highlighted in red). Each point represents mean (n=9) and SEM for OCT4 count. (d) A total of 361 chemistries screened for 24 h included 19 selected monomers printed alone and mixed pairwise (2:1 v/v). (e) OCT4+ hPSC attachment (n=9) was clustered by Euclidean distance measure (intensity scale represents OCT4+ nuclei count) and (f) ranked (high to low) (full list in table S3). Polymer D, (TCDMDA) containing polymers are denoted in red. All letter IDs mentioned are defined in Figure S3.

From this initial screen, nineteen monomers were selected for a second-generation co-polymer screen (denoted in Figure 1c as red letter IDs, and corresponding polymer structure shown in Figure S3). Monomers residing on the y=x line show OCT4 expression in >80% hPSCs, whereas those shifted below the line show decreasing percentage of pluripotency either from loss of expression or attachment of non-viable debris cells. All selected polymers cover a large chemical diversity, show high percentages of OCT4 expression (>80%), but a varied level of attachment. Examples of materials taken forward to the second screen include high attachment, monomer P (tetrahydrofurfuryl acrylate, THFuA; ∼55 ± 35 cells spot^-1^, 93% OCT4 purity) to low attachment monomer S (N-(hydroxymethyl)acrylamide, HMAm; ∼1 ± 2 cells spot^-1^, 100% OCT4 purity). This panel of candidates provided a variety of polymer combinations to be assessed in a second generation screen, where homopolymers were included alone and mixed pairwise as 2:1 v/v (where each monomer was combined as a major and minor component) prior to printing to create a library of 361 chemistries printed in triplicate (Figure 1d).

Screening 361 co-polymers by OCT4+ ReBl-PAT attachment at 24 h by clustering using Euclidean measure distance (Figure 1e) and ranking them by (Figure 1f, see table S3 for polymer names) identified almost 80 chemistries which supported high hPSC attachment (mean cell number > 75 cells spot^-1^, > 90% OCT4+ cells). More than 25% of these polymers contained monomer D, tricylodecane-dimethanol diacrylate (TCDMDA) as a homopolymer and co-polymer (combined pairwise as either a major or minor component) (Figure 1f). TCDMDA also supported attachment to human dental pulp-derived stem cells in previous microarray screening studies as both a homopolymer and co-polymer.^[15]^

To refine the candidate list further, the synergy ratio (SR) of co-polymer combinations was assessed; where cell response to the co-polymer is compared to individual responses of their monomer counterparts (see supplementary information for method) (Figure S4).^[10]^ While most (∼70%) of the 342 combinations were antagonistic or merely additive (SR≤1), ∼30% of the co-polymer features were determined to be synergistic (SR>1) (Figure S4a, b). Consistent with previous studies, synergistic combinations were often related to co-polymers that combined a high attachment monomer with a low attachment monomer .^[10]^ Candidates taken forward for scale-up were chosen on the basis of high cell attachment scores and consideration of the synergistic performance.

The assays above evaluated attachment and retention of pluripotency for 24 h, so our next experiments focused on periods up to 72 h as an indicator of compatibility for longer term expansion in tissue culture plastic (TCP) well-plates (see supplementary information for methods). Seven chemistries (mixed 2:1 v/v) were taken forward for scale-up experiments, including high attachment “candidates” (D:Q, B:L, H:N, D:F) and “controls” for low attachment (E:M, P, B:P; Figure 2a-f, Figure S5). Three of the four high attachment “candidates” applied to 96 well plates (∼3 mm^2^), which evaluated the translation to a ∼100-fold increased surface area compared to a polymer spot (∼0.03 mm^2^), were able to support hPSC attachment and OCT4 pluripotency at 24 h (Figure 2 b-d) and 72 h (Figure 2 e-f) time points. Co-polymer D:Q was found to support high attachment of hPSCs that was most comparable with Matrigel controls, and was thus selected for long-term scale-up experiments.

**Figure 2:**
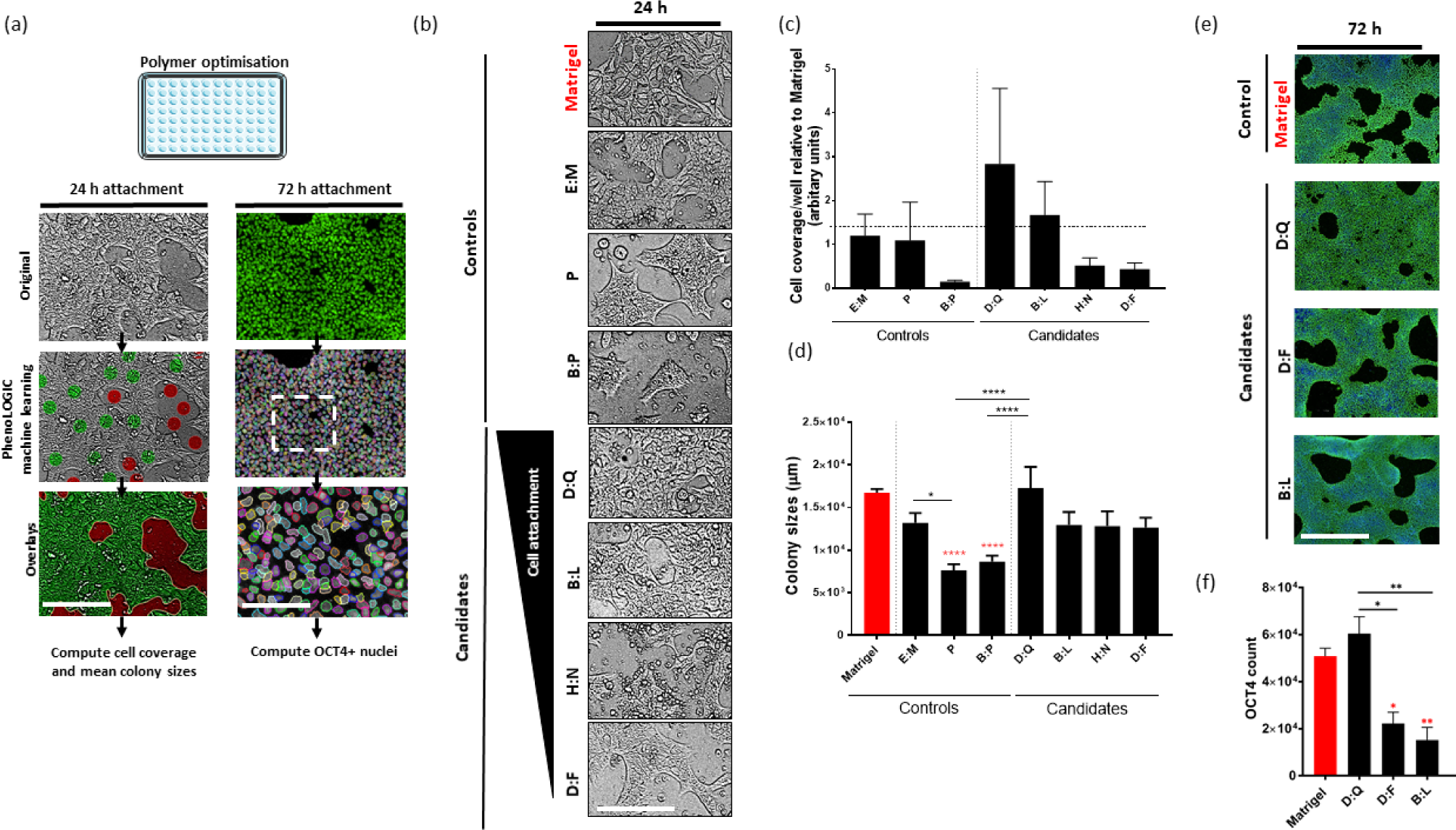
Screening of co-polymer candidates at scale-up. (a) ReBl-PAT hPSCs were cultured in E8 on polymers coated on 96 well plates (n=9/condition). Attachment was assessed at 24 h and 72 h time-points and Matrigel^TM^ was included as a positive control (highlighted in red). All images (5 fields of view/well) were captured (Operetta, Perkin Elmer) and processed using Harmony image analysis software (Perkin Elmer). Cell attachment (cell coverage and mean colony sizes) at 24 h was quantified from brightfield live-cell images processed using scripts developed with PhenoLOGIC machine learning (script training (left centre panel) training: green dots= cells, red dots background, resultant overlays (left bottom panel)) to create a mask for cell coverage/well. At 72 h, hPSCs were fixed and stained for OCT4 expression. OCT4+ nuclei were quantified from fluorescence images (script right panel). (b) Representative brightfield images of ReBl-PATs cultured on polymers (structures to letter IDs in Table S1) and Matrigel^TM^ in E8 medium after 24 h. hPSC attachment on co-polymers are ranked by mean cell coverage/well quantified in (c) relative to Matrigel^TM^ control. (n=9) (d) Mean colony sizes (tightly packed cells with defined outer border) were calculated per field of view (n=15 per condition). Bar graphs represent averaged colony sizes (n) of matrigel (164), E:M (54), P (57), B:P (62), D:Q (54), B:L (53), H:N (47) and D:F (53). One way ANOVA followed by Tukey’s multiple comparison tests (*p<0.05,****p<0.0001) were performed. Statistical differences denoted with red asterisks represent comparisons between polymers and Matrigel. Remaining comparisons are labelled. (e) Representative images and (f) OCT4+ cell attachment per well (n) on polymers at 72 h (∼90% OCT4 expression/ well). One-way ANOVA followed by Tukey’s multiple comparison tests (*p<0.05) (n=9). All graphs represent mean (±SEM). Scale bars = 200μm.

Co-polymer D:Q, which comprises of tricylodecane-dimethanol diacrylate (TCDMDA) mixed with butyl acrylate (BA) (2:1 v/v) was applied to 6-well TCP plates of ∼10cm^2^ base surface area per well (see supplementary information for methods). Surface characterization by TOFSIMS after scaled production confirmed characteristic peaks of TCDMDA (C5H7^+^ m/z = 67.05) and BA (C4H9^+^ m/z = 57.07) (Figure S6). Meanwhile, atomic force microscopy revealed a nanoscale topography in both deformation and modulus images (Figure 3a, b). This indicates that nano scale phase separation of the monomers occurred before polymerization resulting in the equivalent of a blend of poly-BA (minor component, 30% v/v as ∼50nm islands) in a continuous phase of poly-TCDMDA (major component, 70% v/v), rather than a uniform surface that would be representative of a co-polymer. Hence, D:Q will be referred to here as poly(TCDMDA-blend-BA) .

**Figure 3:**
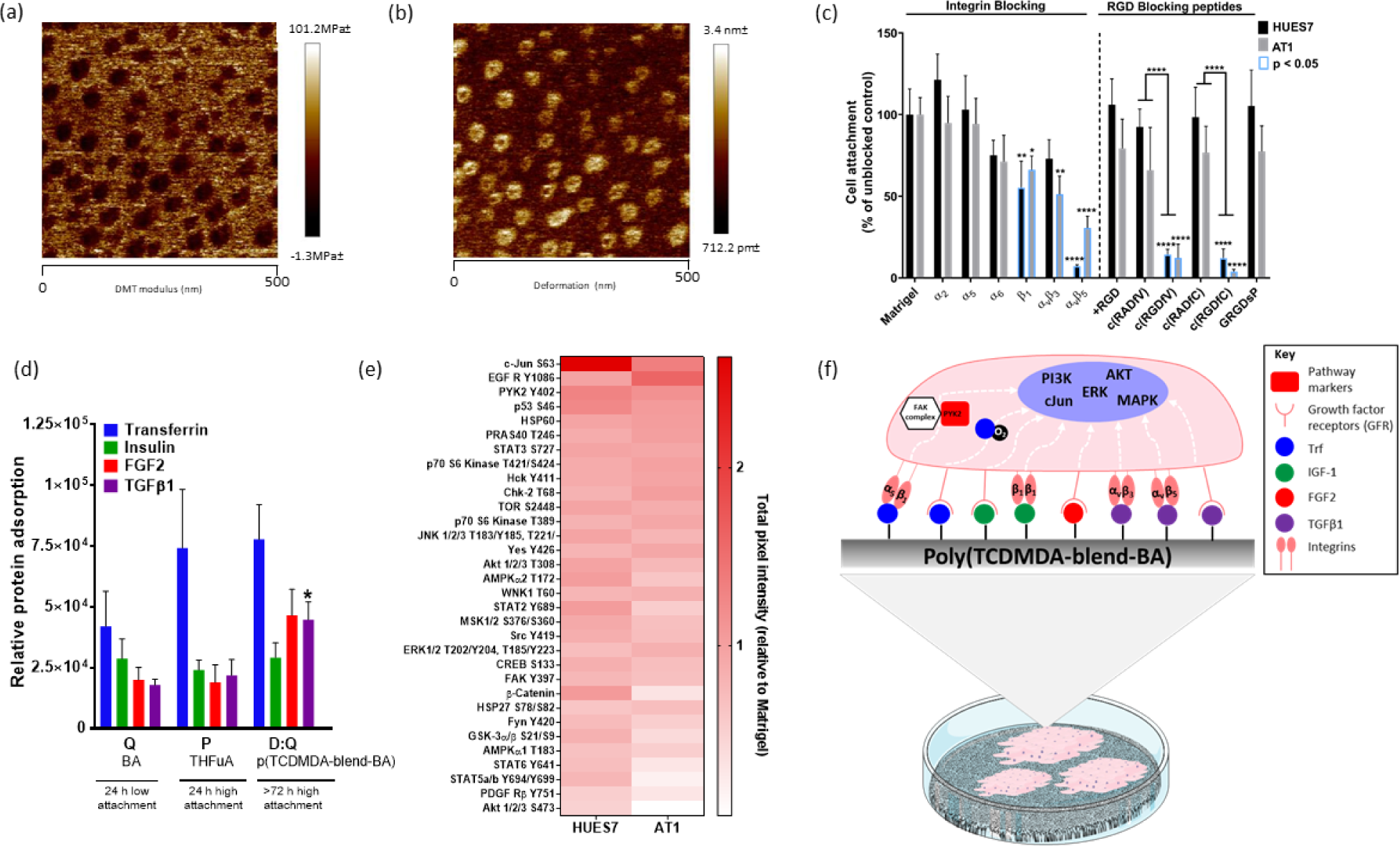
Characterization of poly(TCDMDA-blend-BA) surface. Atomic force microscopy (a) Derjaguin-Muller-Toporov (DMT) modulus and (b) deformation micrographs of poly(TCDMDA-blend-BA) surface coated on poly(styrene) six well plates showing a nanoscale blend of poly-BA (∼50nm islands of minor component, 30% v/v) in poly-TCD (background, major component, 70% v/v). (c) Blocking of integrins and RGD-blocking peptides (see Table S1 for details) for 24 h on poly(TCDMDA-blend-BA) in E8. Bar graphs presented as mean and error bars represent ±STDEV. (N=3, n=3). (d) LESA-MS/MS quantification of adsorbed proteins: FGF2, TGFβ1, insulin and transferrin from E8 medium on polymer surface after 1 h of incubation (N=3). Bar graph represents ±SEM. One-way Anova statistical tests were performed and compared between chemistries for each protein (*p<0.05). (e) Phosphokinase array blots were quantified using Image Studio Software for hESC line HUES7 (N=2, n=4) and hiPSC line AT1 (N=1, n=2). Heatmap represents total intensity values per phosphorylated kinase normalized to background intensity and HSP60 internal control. Graph shows Mean + STDEV. (f) Schematic to summarize identified hPSC and poly(TCDMDA-blend-BA) interactions. The upper panel is a zoomed in image of a single cell-polymer interaction from lower image showing hPSC colonies attached on a well of cultureware coated with poly(TCDMDA-blend-BA). In brief, adsorption of E8 proteins mediate integrin engagement and/or activation of growth factor receptors which subsequently promote key hPSC signaling pathways.

To evaluate whether poly(TCDMDA-blend-BA) could support hPSC attachment, proliferation and expansion across serial passages, we evaluated the performance of ReBl-PAT hiPSCs, and two other lines (AT1,^[14]^ an in-house hiPSC line; HUES-7,^[16]^ a hESC line from Harvard University). At scale-up, we demonstrated that all three hPSC lines cultured on poly(TCDMDA-blend-BA) could readily attach and proliferate, reaching confluency every 72 h across 8 serial passages (Figure S7a, b). The hPSC lines also retained stable karyotypes (46, XY for HUES7; 46, XY for REBl-PAT; 46, XX for AT1; Figure S8, see supporting information for methods) and maintained pluripotent marker expression of OCT4, NANOG, SOX2, TRA181 and SSEA4 confirmed by immunostaining (>80%), flow cytometry (>85%) and quantitative real-time PCR (Figure S9a-c).

We next explored integrin-based mechanisms mediating hPSC attachment on the poly(TCDMDA-blend-BA) surface. To assess this, antibodies and RGD (arginine-glycine-aspartate)-based peptides were used to block integrins known to be highly expressed on the hPSC surface (Figure 3c, see Table S4 for details).^[17, 18]^ Antibody-mediated blocking of vitronectin-binding αvβ3 (AT1, p<0.01) and αvβ5 (p<0.0001) integrins, the Matrigel^TM^ -binding β1 integrin, as well as RGD-peptides binding αvβ3 c(RGDfV), (p<0.0001) and αvβ5 c(RGDfC) (p<0.0001), significantly attenuated hPSC attachment to poly(TCDMDA-blend-BA.^[19–22]^ Expression of key integrins (αv, α5, β1, β4 and β5) was also confirmed through western blotting (Figure S10a, b). Integrin-mediated hPSC attachment to synthetic polymer substrates, notably of β1 and αv integrins, has been previously reported.^[11, 23]^ These data suggest that αvβ3 and αvβ5 heterodimers play an important role for mediating hPSC attachment to poly (TCDMDA-blend-BA), however, integrins are likely to interact in a complex manner and therefore can form a number of different homo- or hetero-dimers.

Having explored the adhesion molecules expressed by the hPSCs, we next evaluated protein absorption to the chemically defined culture substrate. Adsorption of proteins from E8 culture medium on poly(TCDMDA-blend-BA) critical to promoting integrin mediated hPSC attachment were identified by liquid extraction surface analysis-tandem mass spectrometry (LESA-MS/MS).^[24]^ Adsorption of medium-derived proteins to the polymer interface was assessed in the absence of cells, using a rinsing procedure designed to retain species bound to the surface for analysis by surface sampling (see supplementary information for methods). Whilst minimal in its formulation, E8 includes key pluripotency-maintaining factors: basic fibroblast growth factor (FGF2) and transforming growth factor beta 1 (TGF-β1); as well as insulin and transferrin, which are required for hPSC attachment and survival.^[1]^

The amount of each protein adsorbed by the polymer poly(TCDMDA-blend-BA) capable of supporting attachment for >72 h was compared to low and high attachment polymers from the short-term 24 h attachment experiments. These are denoted as monomers Q and P in Figure 1c, 2, polyBA and polyTHFuA respectively in Figure 3d. Of the four proteins detected, significantly higher TGFβ1 adsorption was observed on poly(TCDMDA-blend-BA) (p<0.05) compared to polymers which only supported attachment for 24 h. The αvβ3 and αvβ5 integrin heterodimers have been reported to induce extracellular matrix-bound TGFβ1 release and activation,^[25]^ with both integrin heterodimers being shown in this study to mediate hPSC attachment to poly(TCDMDA-blend-BA) (Figure 3c). Since TGFβ1 is an important factor for maintaining hPSC pluripotency, it is likely that TGFβ-binding contributes to the success of poly(TCDMDA-blend-BA) in long-term maintenance of hPSCs in their undifferentiated state.

Intracellular signaling profiling of hPSCs cultured on poly(TCDMDA-blend-BA) confirmed maintenance of stem cell integrity. We now wished to explore the impact of these interactions on molecular signaling pathways. A phospho-kinase array kit (R&D) containing a panel of critical markers was used to assess intracellular signaling events in two independent hPSC lines (AT-1, HUES7) (Figure 3e).

Expression profiles for several kinases appeared to be cell line specific. However kinases: c-Jun (S63), EGFR (epidermal growth factor receptor, Y1086) and PYK2 (tyrosine kinase encoded by PTK2B, Y402) were differentially phosphorylated on poly(TCDMDA-blend-BA) compared to Matrigel^TM^ in both hPSC lines (Figure 3e). These kinases activate important downstream hPSC proliferation pathways (including phosphoinositide 3-kinase (PI3K) / AKT, c-Jun, extracellular-signal-regulated kinase (ERK)/ mitogen-activated protein kinases (MAPK)), most likely initiated by integrin engagement (including αv, β1 and β3) based on the integrin-interface characterization experiments on poly(TCDMDA-blend-BA) (Figure 3c and Figure S10).^[26–28]^ Alternatively, adsorbed E8 factors, all four of which were confirmed for poly(TCDMDA-blend-BA), can promote hPSC signaling pathways by activating important growth factor receptors (eg. insulin-like growth factor receptor (IGFR), epidermal growth factor receptor (EGFR), fibroblast growth factor receptor (FGFR) and transferrin receptor (TrFR)) for hPSC signaling (proposed mechanism summarized in Figure 3f).

In addition to understanding the mechanisms of hPSC attachment and signaling, the value of these pluripotent cells is differentiation to multiple lineages for use in biomedical application and regenerative medicine. Therefore, after serial passaging on poly(TCDMDA-blend-BA), directed differentiation of hPSCs towards the three germ layers was performed.^[11]^

Definitive endoderm SOX17 and FOXA2 positive cells were achieved after two days of WNT pathway activation with the GSK-3 inhibitor CHI99021 (Figure S11a). hPSC differentiation to SOX1 and PAX6 positive neural progenitors of the ectoderm lineage was achieved with modulators of the transforming growth factor beta (TGF-β) superfamily (dual SMAD inhibitors dorsomorphin and SB431542) and WNT (XAV393) pathways (Figure S11b). Functional contractile cardiomyocytes (mesoderm) cells were formed using modulators of the TGF-β (activin A and BMP4) and WNT (KY02111 and XAV393) pathways for 8-12 days (Figure S11c). Thus, achieving multi-lineage differentiation, confirmed that stem cell integrity was retained after serial passaging on poly(TCDMDA-blend-BA).

In summary, a high throughput combinatorial materials discovery approach identified poly(TCDMDA-blend-BA) for long-term hPSC culture in commercial defined E8 medium, without the need for addition of xenogeneic factors or attachment-mediating proteins from culture medium. This simplified system has advanced our understanding of the factors at the bio-interface between media and polymer controlling stem cell response. Protein adsorption of chemically defined E8 medium was compared between different polymers using mass spectrometry methods. Together with cell-based assays, which investigated cell-polymer interactions, this enabled us to report on mechanisms for cell adhesion and intracellular signaling which corroborated well with our previous work and other culture systems used routinely within the field. ^[1, 5–8, 10, 11, 13, 19, 22, 23]^ This material constitutes a break-through for the application of hPSCs, addressing current limitations of scalability and cost in the production of cells for routine clinical-scale production.

## Experimental Section

Complete methodology of polymer synthesis, characterization and cell-based assays can be found in the Supporting information.

## Supporting Information

Supporting Information is available from the Wiley Online Library or from the corresponding author(s).

## Acknowledgements

J.T, A.N and L.B contributed equally to this work. This work was supported by the Engineering and Physical Sciences Research Council Program Grant in Next Generation Biomaterials Discovery [EP/N006615/1], as well as Wellcome Trust [201457/Z/16/Z], British Heart Foundation (BHF) [grant numbers: SP/15/9/31605, PG/14/59/31000, RG/14/1/30588, RM/13/30157, P47352/CRM], National Centre for the Replacement, Refinement, and Reduction of Animals in Research (NC3Rs) [CRACK-IT:35911–259146, NC/K000225/1, NC/S001808/1]. The authors would also like to acknowledge Dr. Christopher Carroll (Biodiscovery Institute, University of Nottingham) for his technical assistance with processing proteome profiling arrays.

## Conflict of interest

The authors declare no conflict of interest

## Supporting Information

### Routine cell culture

Three hPSC lines used in this study, including the hESC line, HUES7 and the hiPSC cell lines: ReBl-PAT derived from a skin punch biopsy from a male subject and AT1 derived from dental pulp of a female subject (as previously described), ^[1]^ were routinely maintained on 1:100 Matrigel coating (BD Biosciences, UK) in Essential 8^TM^ medium (E8, LifeTechnologies). Cells were passaged at 70-80% confluency by washing once PBS, followed by incubation with TrypLE Select (LifeTechnologies) for 3 minutes at 37°C, with tapping of flasks to dissociate cells.

### Polymer microarray synthesis and preparation

Polymer microarrays were fabricated using methods previously described.^[2, 3]^ Briefly, polymer microarrays were printed onto polyHEMA (4% w/v Sigma, in ethanol (95% v/v in water)) dip coated glass slides using a XYZ3200 dispensing station (Biodot) and quilled metal pins (946MP6B, Arrayit) under an argon atmosphere maintaining O2 ˂ 2000 ppm, 25°C and 35% humidity. Polymerization solutions consisted of monomer (50% v/v) in dimethylformamide with photoinitiator 2,2-dimethoxy-2-phenyl acetophenone (1% w/v), and were polymerized in-situ using UV light irradiation. Three replicates of 284 monomers were printed per slide for the first generation array (see Figure S1 for structures and Table S1 for monomer list). For the second generation array, the polymerisation solutions consisted of major and minor monomers in a 2:1 (v/v) ratio. Three replicates of 342 co-polymers combinations were printed per slide. Monomers were purchased from Sigma, Scientific Polymers and Polysciences and were used as received. Top and bottom array surfaces were sterilised with UV light for 15 minutes and washed with sterile Ca^2+^/Mg^2+^ -free Phosphate Buffer Saline (PBS, Gibco) before culturing with hPSCs.

### Microarray screening and data acquisition

0.75x10^6^ REBl-PAT cells were seeded in E8 medium supplemented with 10μM Y-27632 (ROCKi, Tocris Bioscience) on each array and incubated at 37°C with 5% CO2 for 24 h and 48 h timepoints at which point array samples were fixed with 4% paraformaldehyde for quantification. Arrays were immunostained for OCT4 expression and counterstained with 4’,6-diamidino-2-phenylindole (DAPI) (see immunostaining methods for full details) before being mounted with Vectashield Antifade mounting medium (Vector Laboratories and imaged using automated fluorescence microscopy (IMSTAR). Attachment was analysed in CellProfiler ver. 2.2.0 (Broad Institute).^[4]^ Manual background correction was also applied to images prior to using in-built “identify primary objects” algorithm using a three-class Otsu adaptive thresholding method to identify and quantify nuclei in DAPI and OCT4 channels with manual check for quality control. Assessment of co-polymer combinations for second generation screen can be readily performed using a synergy ratio (SR). Taking the response of major (y1) and minor (y2) monomers alone, normalised to the fraction (m) present in the co-polymer (y12), the SR can be calculated using the equation: 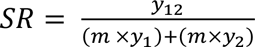

A synergistic combination, SR>1, indicates that the cell response for the co-polymer is greater than the response of the individual monomers. Whilst an additive/counteractive combination, SR<1, indicates that the cell response for the co-polymers is less than the response of the individual monomers.

### hPSC assessment of polymer candidates coated on 96-well plates

ReBl-PAT hPSCs were seeded at 4.5x10^4^ cells/cm^2^ on co-polymers selected for scale-up in E8 medium supplemented with Y-27632 where each co-polymer was tested in triplicate wells. Matrigel controls were also included for comparison. Images of five separate fields were obtained per well (n=3 independent repeat) using the Operetta high-content imaging system (Perkin Elmer). Images were analysed using Harmony high-content image analysis software (Perkin Elmer) developed with PhenoLOGIC machine learning algorithms to quantify percentage cell coverage (relative to total areas imaged per well) and mean area of colonies (total cell coverage/no. of colonies). Adhered cells at 72 h were fixed in 4% paraformaldehyde and immunostained for OCT4 and fluorescence microscopy using the Operetta and Harmony was used to quantify total and OCT4+ nuclei (5 fields/well).

### Production of polymer coated 6-well plates

Monomers for polymerisation, consisting of individual monomers or 2 monomers mixed at 2:1 (v/v), were mixed in a 9:1 (v:v) ratio with a 10 wt % solution of photoinitiator 2,2-dimethoxy-2-phenyl acetophenone in isopropyl alcohol and coated onto oxygen plasma treated (pi=0.09 mbar, 100 W, 13.56 MHz RF generator for 10 minutes) tissue culture plastic well-plates. These were then polymerised by exposure to UV light (365 nm, 2 x 15 W, 10 cm distance) for 1 h in an argon glovebox (<2000 ppm O2). After polymerization, well-plates were washed three times with isopropanol to remove unreacted polymer, and soaked in dH2O for 48 h at 37°C. Well-plates were subsequently sterilized with 70% IMS and washed three times with sterile PBS before use.

### Surface chemistry analysis

The surface chemistry of array slides and 6-well plates was assessed using time-of flight secondary ion mass spectrometry (ToF-SIMS) and atomic force microscopy (AFM).

### ToFSIMS

Measurements were taken using a TOF-SIMS 4 (IONTOF GmbH) instrument using a 25kV Bi3^++^ primary ion source with a pulsed target current of ∼1pA as previously described.^2^

### AFM

Hydrated AFM measurements were acquired using a Bruker Dimension FastScan in PeakForce^TM^ mode using SCANASYST-FLUID+ probes. Samples assessed for surface analysis were incubated in ultrapure MilliQ water (18.2 Ohm) and the probes were calibrated using a 2.6 GPa Bruker polystyrene film sample.^[2]^

### Protein adsorption analysis

Sterilized and washed polymer coated 6 well-plates were incubated in E8 medium supplemented with 10μM Y-27632 dihydrochloride for 1 h at 37°C. Plates were washed with dH2O (18.2 MΩ, ElgaPure LabWater). Proteins were digested in-situ using microwave-assisted techniques using 0.05 μg/µL trypsin (sequencing grade; Promega, UK) in 100mM ammonium bicarbonate (BioUltra,≥99.5%, Sigma-Aldrich) adapted from previously described methods.^[5]^ Standard methods were used to extract proteins using an extraction solution consisting of acetonitrile (CHROMASOLV®, Riedel-de Haen) and 200 mM ammonium acetate (≥99.0%; Sigma-Aldrich, Gillingham, UK) (1:9 v/v) in LC-MS grade water (CHROMASOLV®, Riedel-de Haen). Samples were analysed by liquid extraction surface analysis-mass spectrometry (LESA-MS) and introduced to a TriVersa Nanomate (Advion Biosciences, Ithaca, NY) coupled to a Q Exactive plus mass spectrometer (Thermo, San Jose, CA) via nanoelectrospray ionisation (ESI Chip™, Advion Biosciences) using 1.6 kV voltage and 0.6 psi gas pressure (N2).

### hPSC serial passaging on polymer coated 6-well plates

hPSCs were seeded at 7x10^4^ cells/cm^2^ in E8 medium supplemented with 10μM Y-27632 dihydrochloride for the initial 24 h of culture. Medium was exchanged every 24 h until cells reached 70-80% confluency at 72 h when cells were fixed or passaged by dissociating with TryPLE select (as described above). hPSCs growth was assessed using an automated cell-viability counter (CEDEX Hi Res Analyser) at each passage (every 72 h). Doubling time (www.atcc.org; [duration of culture x log2] / [log10 (final cell concentration/seeding concentration)] was calculated for hPSCs and was plotted cumulatively. After 5 serial passages HPSC were karyotyped as previously described.^[1]^

### Flow cytometry

hPSCs serially passaged on polymer substrate (≥ 3 passages) were dissociated into single-cell suspension and fixed with 4% paraformaldehyde. Samples were permeabilized with 0.1% Tween-20 in PBS for intracellular markers and incubated with primary antibodies NANOG (1:100, APCH7 conjugated, BD Biosciences, 560109), SOX2 (1:20, Alexa Fluor 647-conjugated, R&D Systems, IC2018R), TRA181 (1:100, PE-conjugated, Invitrogen, 12-8883-82) and SSEA4 (1:20, fluorescein-conjugated, R&D Systems, FAB1435F) diluted in PBS for 1hr at RT. The FC500 flow cytometer (Beckman Coulter) was used to acquire measurements and expression was quantified with Kaluza analysis software (Beckman Coulter).

### Attachment blocking

hPSCs were harvested and re-seeded in E8 medium with the addition of integrin blocking antibodies (10μg/ml for each antibody) or RGD-blocking peptides (15μg/ml) for 24 h (see Table S2). Cells were washed three times with PBS, fixed with 4% paraformaldehyde and counterstained with DAPI. Fluorescence images acquired using the Operetta (Perkin Elmer) were quantified for total nuclei count per condition in Harmony image analysis software (Perkin Elmer).

### Integrin expression by Western Blot

hPSCs serially passaged on polymer (≥ 3 passages) were lysed using RIPA buffer (Cell Signalling Technologies #9806) supplemented with PMSF (Phenylmethylsulfonyl fluoride, Sigma 10837091001). Total lysate protein was determined using Pierce BCA Protein Assay Kit (Thermo Fisher Scientific # 23225) following manufacturer’s instructions. LDS NuPAGE Sample Buffer (4X) with 2.5% 2-mercaptoethanol was added to 30μg of protein lysate and run on NuPAGE NOVEX Bis-Tris Gels with MOPS SDS Running Buffer (Thermo Fisher Scientific #NP0008, #NP0001). Samples were transferred to an Amersham Protran 0.45m nitrocellulose blotting membrane (GE Healthcare Life Science #10600124). Membranes were incubated with following antibodies α5 (#4705), αv (#4711), β1 (#9699), β4 (#14803) and β5 (#3629)integrins (all purchased from Cell Signalling Technology and diluted 1:500), Nanog (clone 7F7.1, Millipore, MABD24, 1:500) and β-actin (Millipore, MA1140, 1:2000). Membranes were developed using West Pico PLUS Chemiluminescent Substrate (Thermo Fisher Scientific #34577) on an LAS-400 Imaging system.

### Proteome Profiler Array

Human Phospho-Kinase Array (R&D systems, ARY003B) was performed according to manufacturer’s instructions (www.rndsystems.com) on hPSCs serially passaged on polymer and Matrigel^TM^ in parallel (≥ 3 passages). Array blots were imaged using ImageQuant LAS-4000 (Fujitsu Life Sciences) and analysed using Image Studio Software (LI-COR, version 5.2.5) where individual total signal intensity was measured by manual gating. All intensity values were normalized to background intensity and HSP60 internal control. Changes were quantified by comparison between Matrigel^TM^ and polymer conditions.

### Tri-lineage differentiation

hPSCs serially passaged (≥ 3 passages) were harvested and seeded at 2x10^4^ – 1x10^5^ cell/cm^2^ and expanded in E8 medium for 2 days with daily media exchanges. All directed differentiation protocols were performed on hPSCs at day 2. For definitive endoderm differentiation, media was replaced by RPMI supplemented with B27 without insulin (LifeTechnologies 0080085-SA) and CHIR99021 (2μM; STEMCELL Technologies, 72052) for a further 2 days with daily media exchanges. To produce neural progenitors of the ectoderm lineage, media was replaced by Advanced DMEM/F-12 (LifeTechnologies) supplemented with 1% L-glutamine (Life Technologies), 1% CD Lipid Concentrate (Life Technologies) 7.5μg/ml Transferrin (Sigma-Aldrich), 14μg/ml Insulin (Sigma Aldrich), 0.1mM β-mercapto-ethanol, 10μM SB431542 (Tocris) and 1μM Dorsomorphin-1 (Tocris) and 2μM XAV939 (STEMCELL Technologies) for 5 days with daily media exchanges. Differentiation to cardiomyocytes was achieved using methods previously described.^[1]^

### Immunostaining

Adherent cells were fixed in 4% paraformaldehyde (Sigma-Aldrich, UK) at room temperature (RT) for 20 minutes and permeabilized with 0.1% Triton-X100 (Sigma-Aldrich, UK) in PBS at RT for 15 minutes. Non-specific binding was blocked with 4% serum (Sigma-Aldrich, UK) in PBS at RT for 1 h. Samples were incubated overnight at 4°C with primary antibodies OCT4 (1:200, Santa Cruz Biotechnology, SC-5279), TRA181 (1:200, Millipore, MAB4381), SSEA4 (1:100, Millipore), FOXA2 (1:500, Sigma-Aldrich 07-633), SOX17 (1:100, R&D AF1924), SOX1 (1:100, R&D AF3369), PAX6 (1:100, R&D AF8150) and cardiac α-actinin (1:800, Sigma-Aldrich A7811) diluted in blocking solution with the addition of 0.1% Triton X-100 for nuclear stains. Samples were washed with 0.1% Tween-20 (Sigma-Aldrich, UK) and incubated with Alexa Fluor secondary antibodies (Life Technologies) 1:400 in blocking solution for 1 h at RT in the dark. Cells were washed with 0.1% Tween-20 and nuclei were counterstained with 0.5μg/ml DAPI (4’,6-diamidino-2-phenylindole, Sigma-Aldrich D9542).

### Statistical tests

Experiments were performed in at least three independent experiments unless otherwise stated. Statistical tests (as stated in text) were performed using GraphPad Prism (version 8.1.2, San Diego CA). Heatmaps were plotted using the heatmap.2 function from the gplots package version 3.1.0.2 in combination with the RColorBrewer package version 1.1-2. Clustering and dendrograms for heatmaps were produced using the complete method with Euclidean distance measure.^[6]^

---

**Figure S1:**
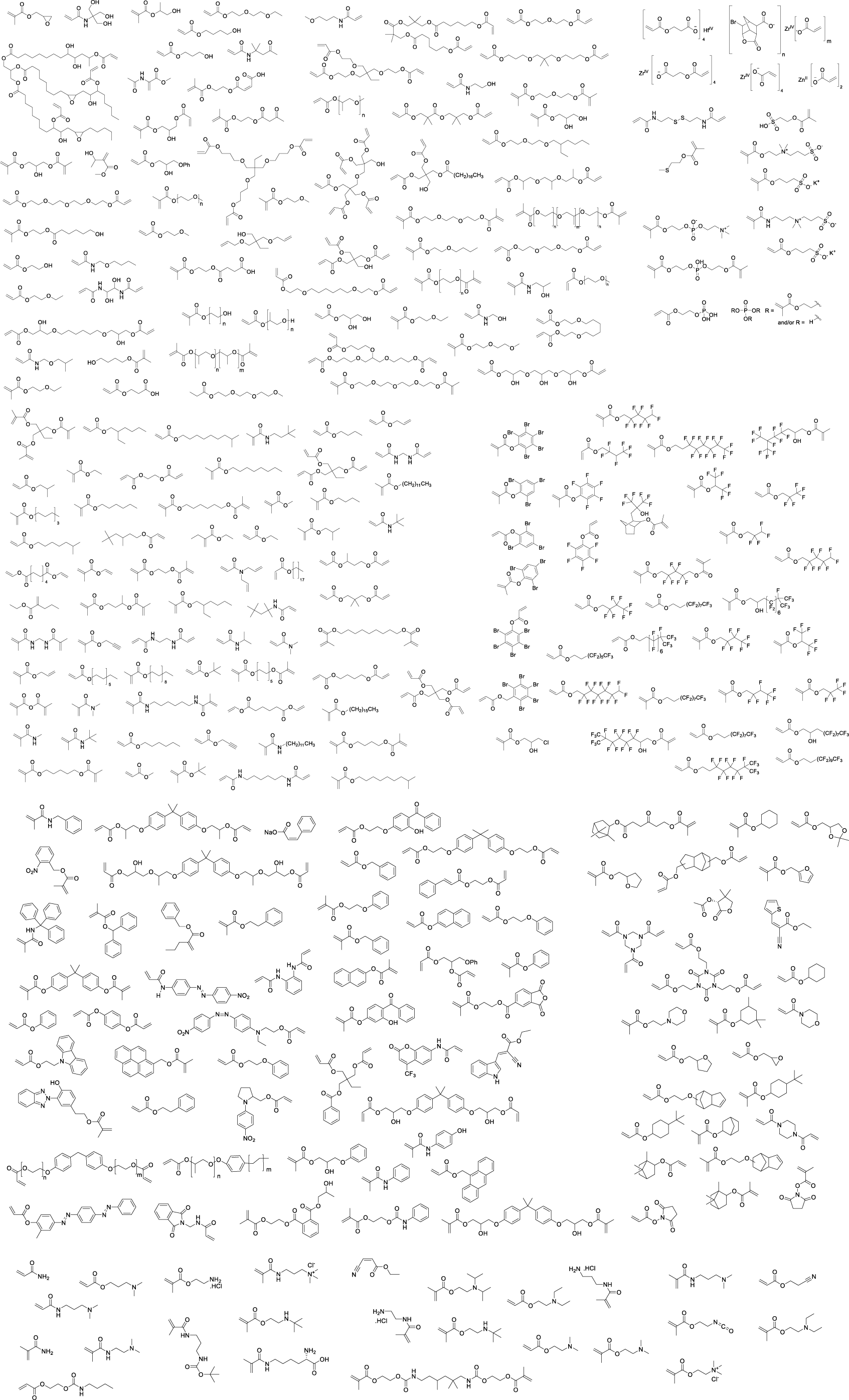
Monomer structures of 284 materials used for the first generation microarray screen.

**Figure S2.**
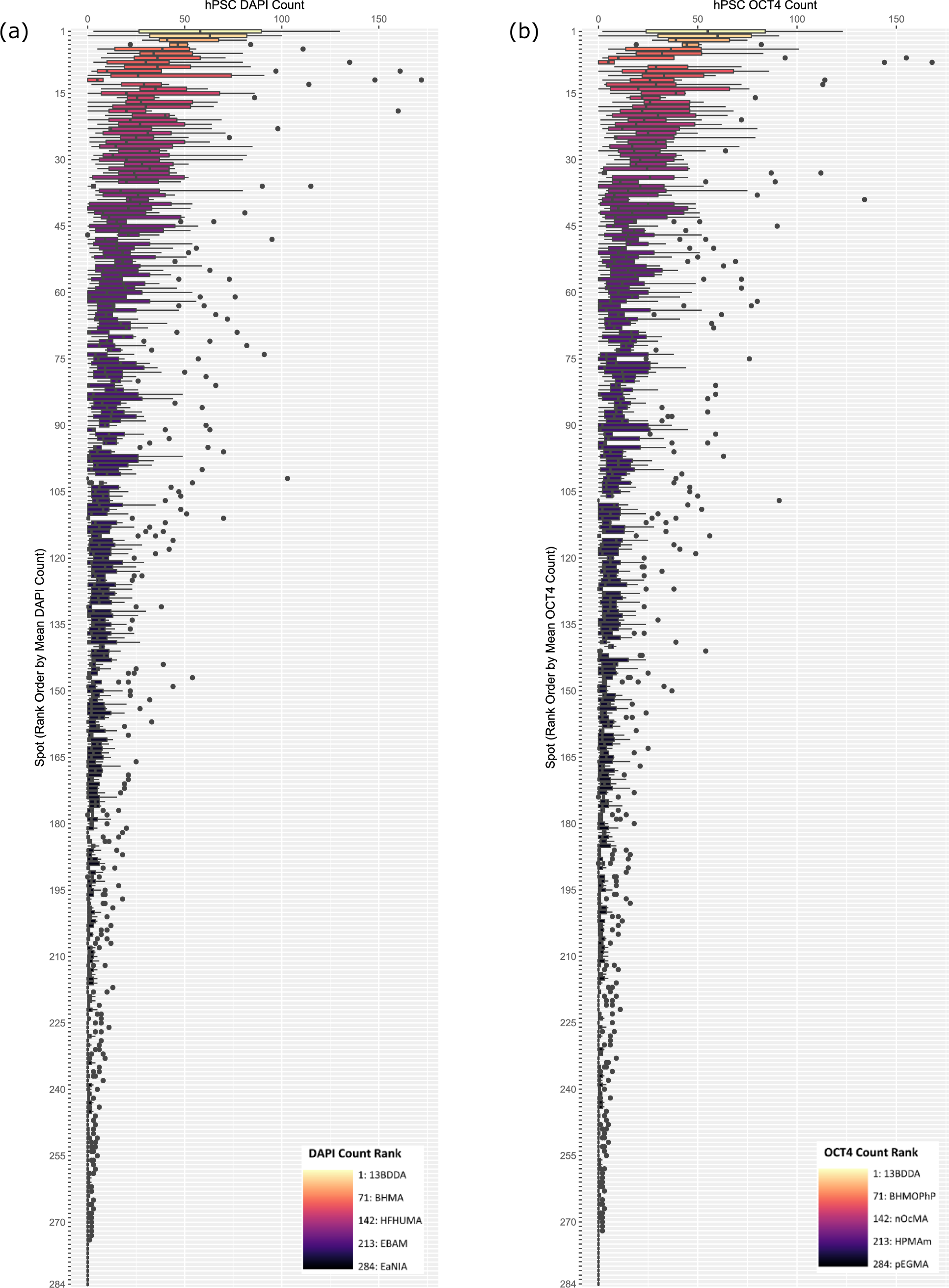
(a) Total cell number (DAPI count) (b) and OCT4+ count of REBl-PAT hPSCs on 284 monomer microarray ranked high to low (denoted light to dark, see legend) after 24 h. (See Table S2 for rank order 1-284)

**Figure S3:**
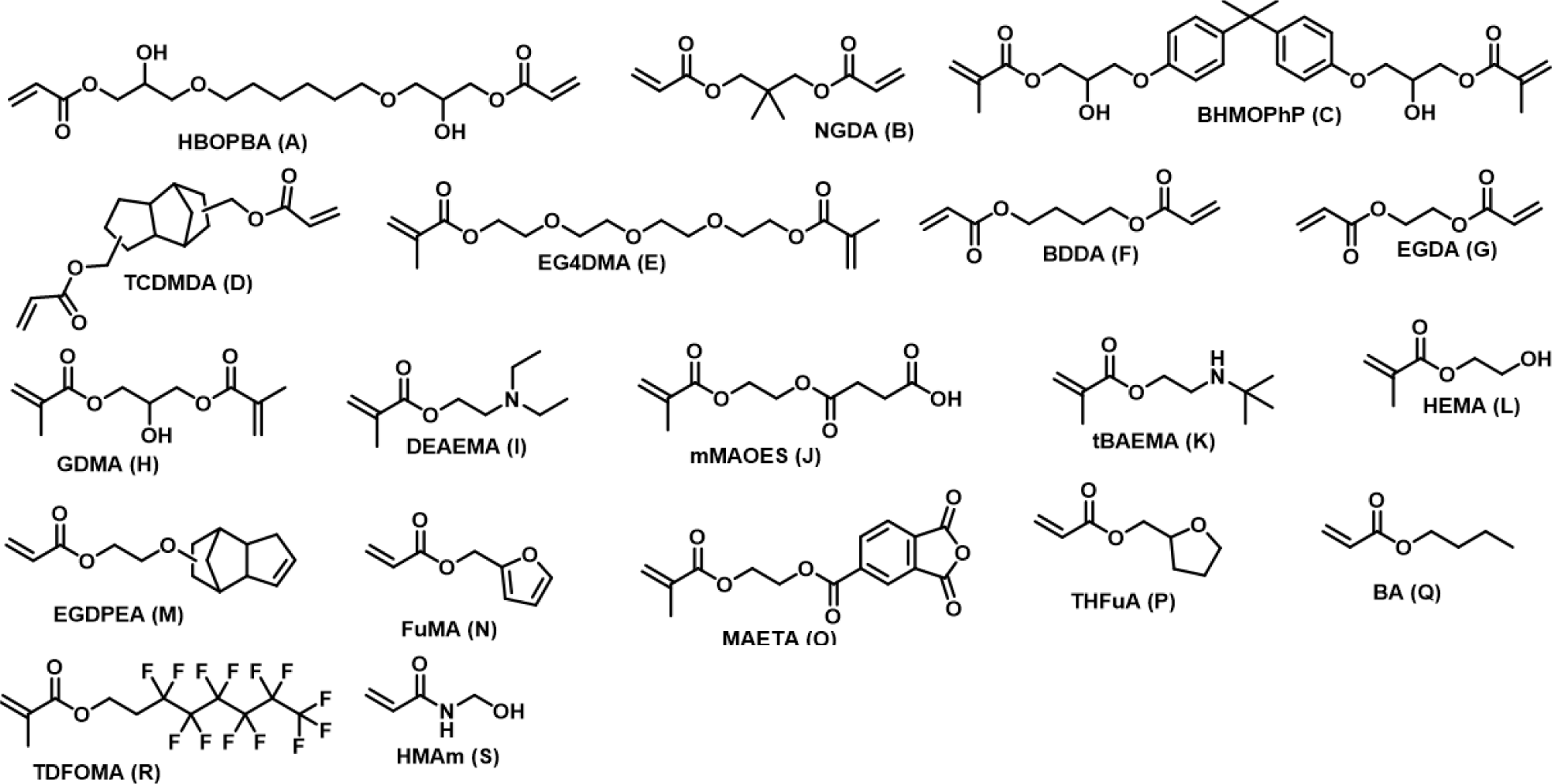
Monomer structures of 19 materials selected for second generation co-polymer screen labelled A-S as referred to in main text.

**Figure S4:**
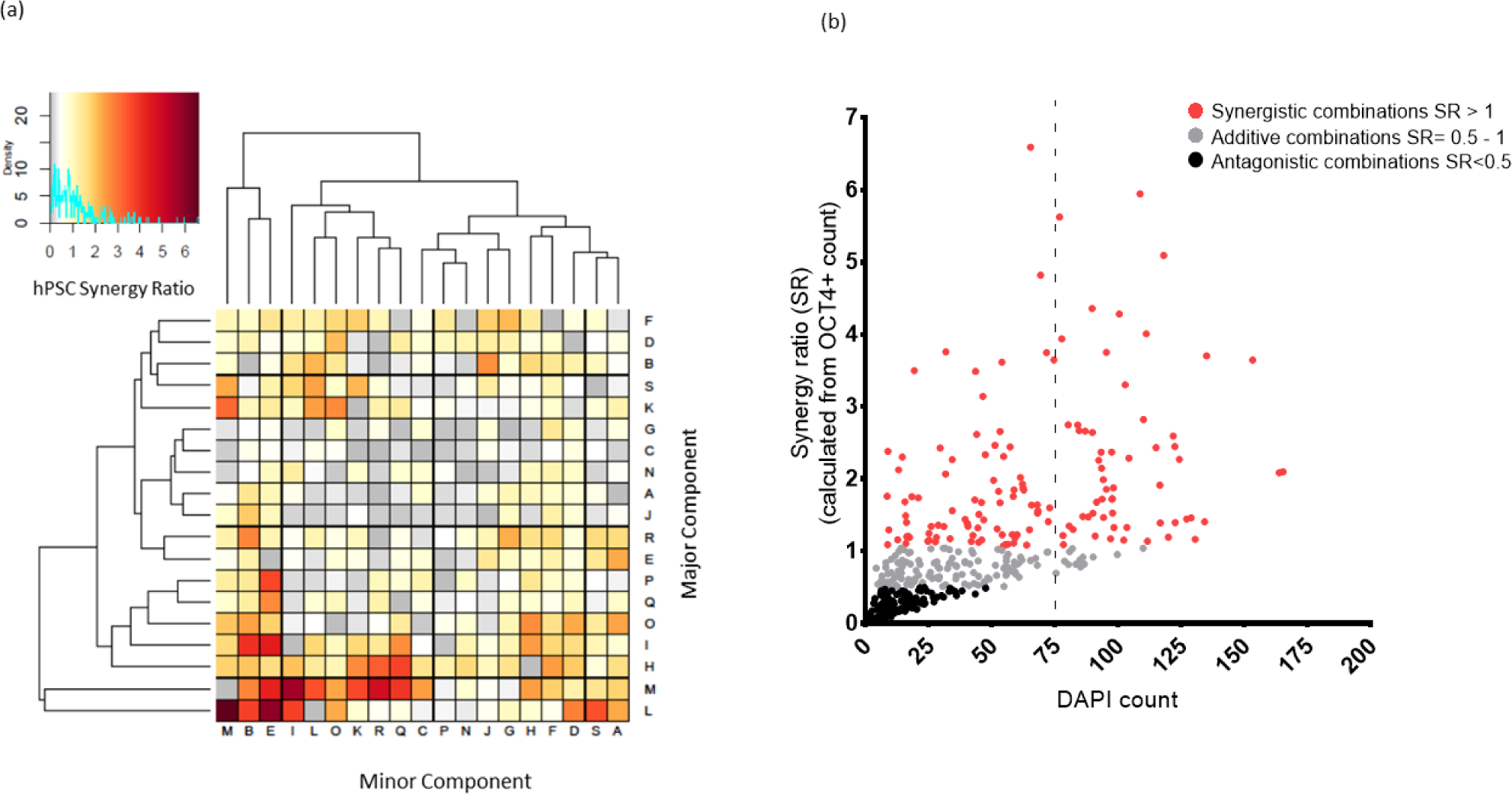
(a) Synergy of co-polymer combinations were quantified as a ratio of OCT4+ attachment for co-polymer to their corresponding homopolymer components (see supplementary information for methods) clustered by Euclidean distance measure. Synergy ratios (SR) >1 are synergistic combinations (denoted yellow -red), SR values = 1 are additive combinations (denoted in white) and SR values <1 are antagonistic combinations (denoted in grey). All letter IDs mentioned are defined in Figure S2. (b) SR scores were plotted against average total cell number (n=9, where n represents the no. of polymer spots). Data has been defined as synergistic (red), additive (grey) or antagonistic. Data points to the right of dotted line represent high attachment polymers. All attachment data is summarized in table S3.

**Figure S5:**
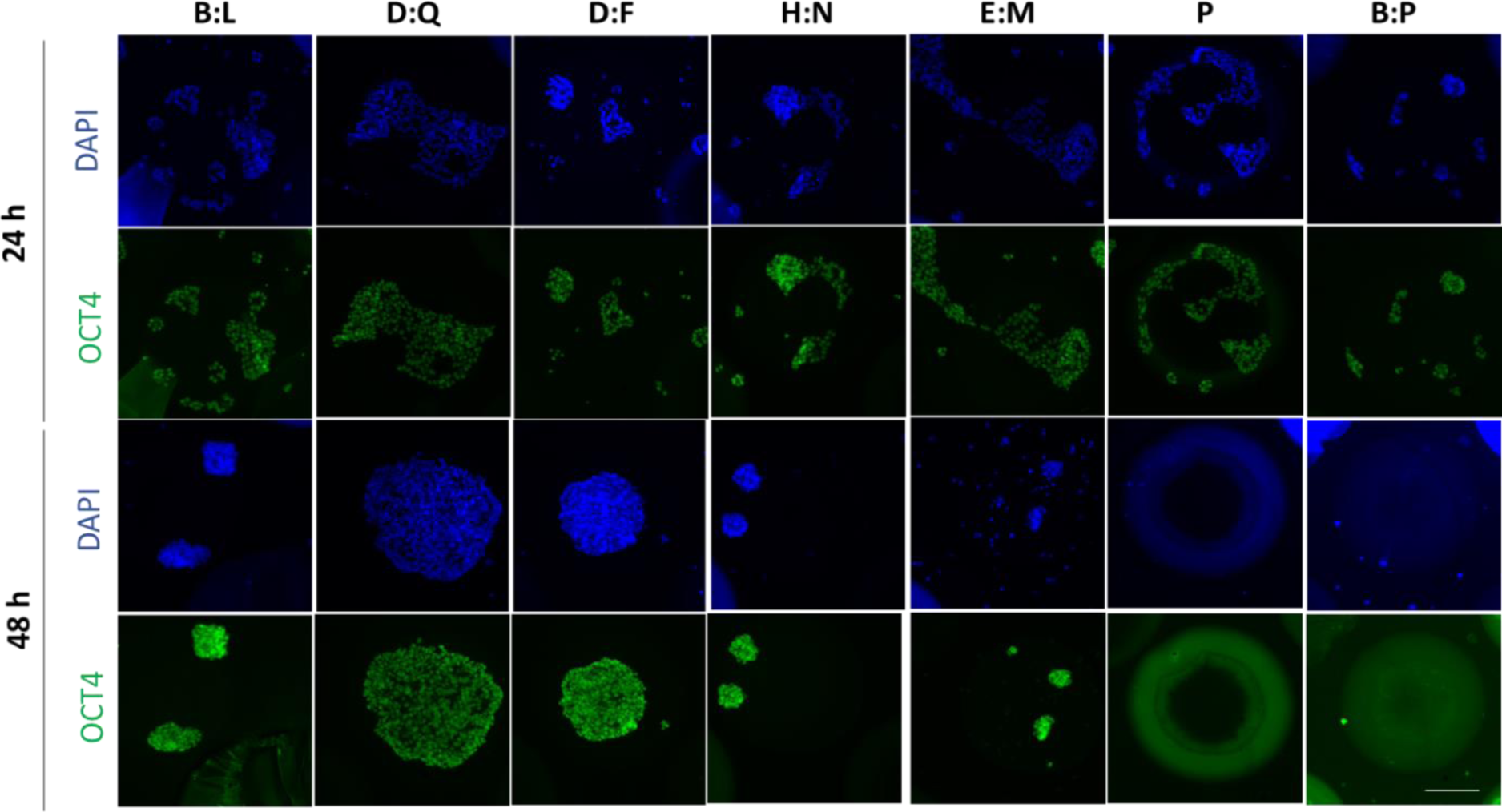
Representative images of OCT4 and DAPI stained REBl-PAT attachment on candidate polymers for scale-up on second generation polymer arrayed slides seeded at 0.75 x10^6^ cells/ array at 24 h and 48 h time points. See Figure S3 for polymer IDs. Scale bar represents 100μm.

**Figure S6:**
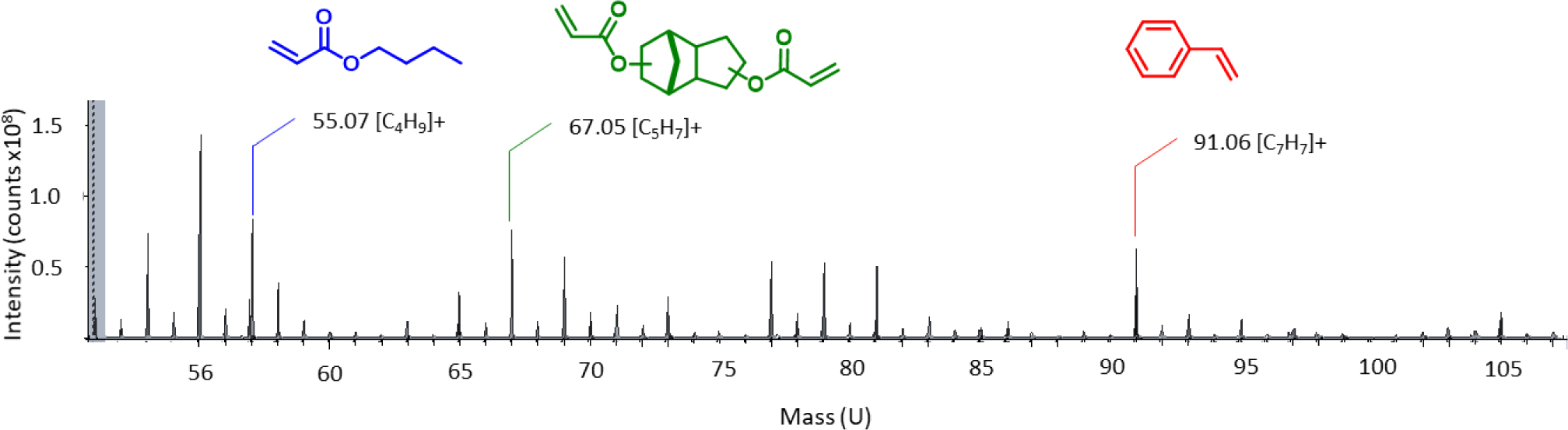
TOFSIMS analysis of poly(TCDMDA-blend-BA) surface on poly(styrene) based tissue culture six well-plates. Ions characteristic of polyBA ([C4H9]^+^ m/z = 57.07, polyTCDMDA ([C5H7]^+^ m/z = 67.05) and poly(styrene) ([C2H7]^+^ m/z = 91.06).(N=3, area analysed =3x3mm, constituent monomers shown for references)

**Figure S7:**
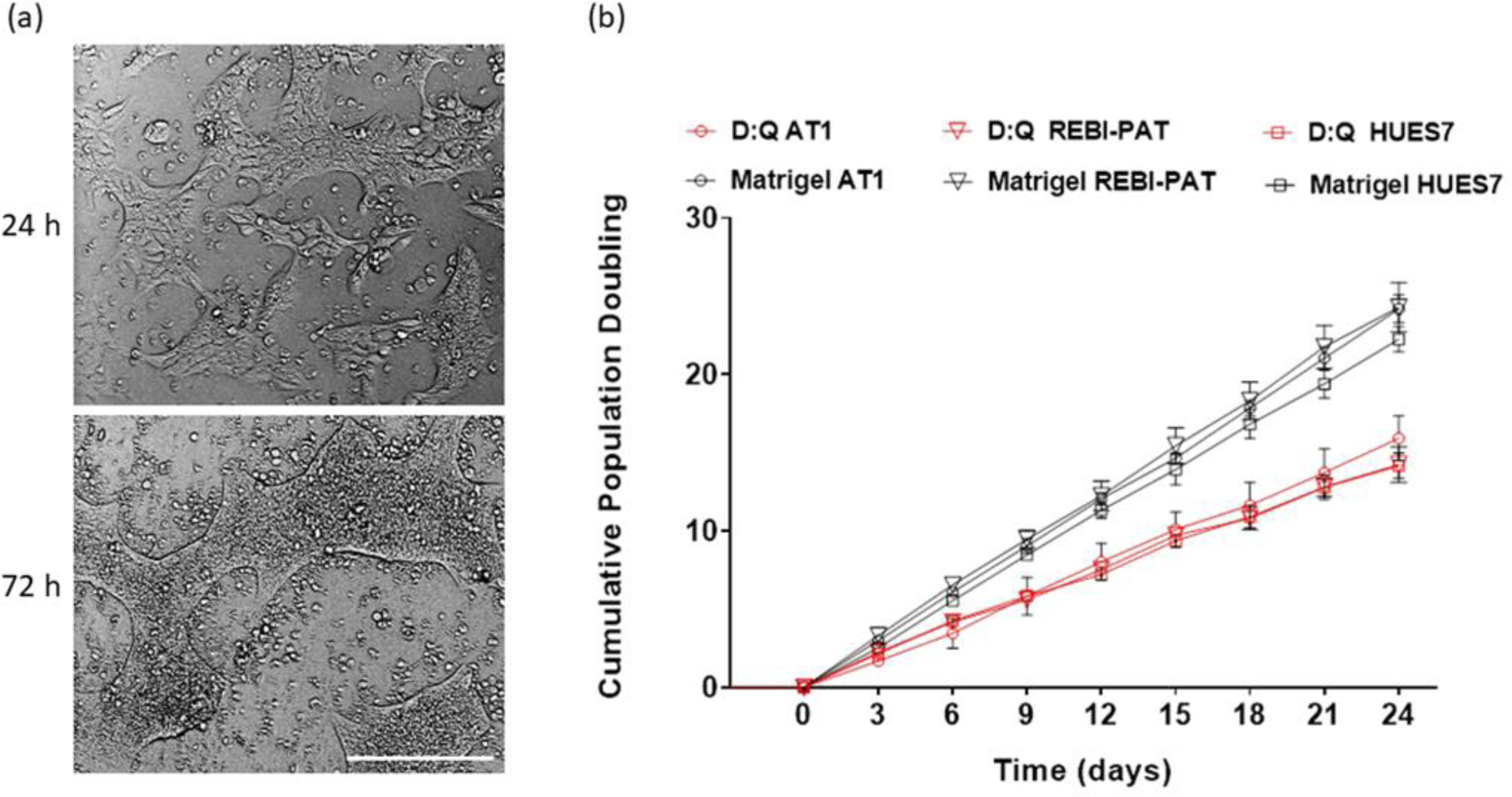
Scaled-up hPSC culture on D:Q (poly(TCDMDA-blend-BA)) (a) Representative brightfield images of ReBl-PAT attachment and maintenance at 24 h and 72 h of culture. (b) Growth curves of hESC: HUES7 and hiPSC AT1 and ReBl-PAT lines on poly(TCDMDA-blend-BA) and Matrigel^TM^ presented as cumulative doubling time (for equation see methods).

**Figure S8:**
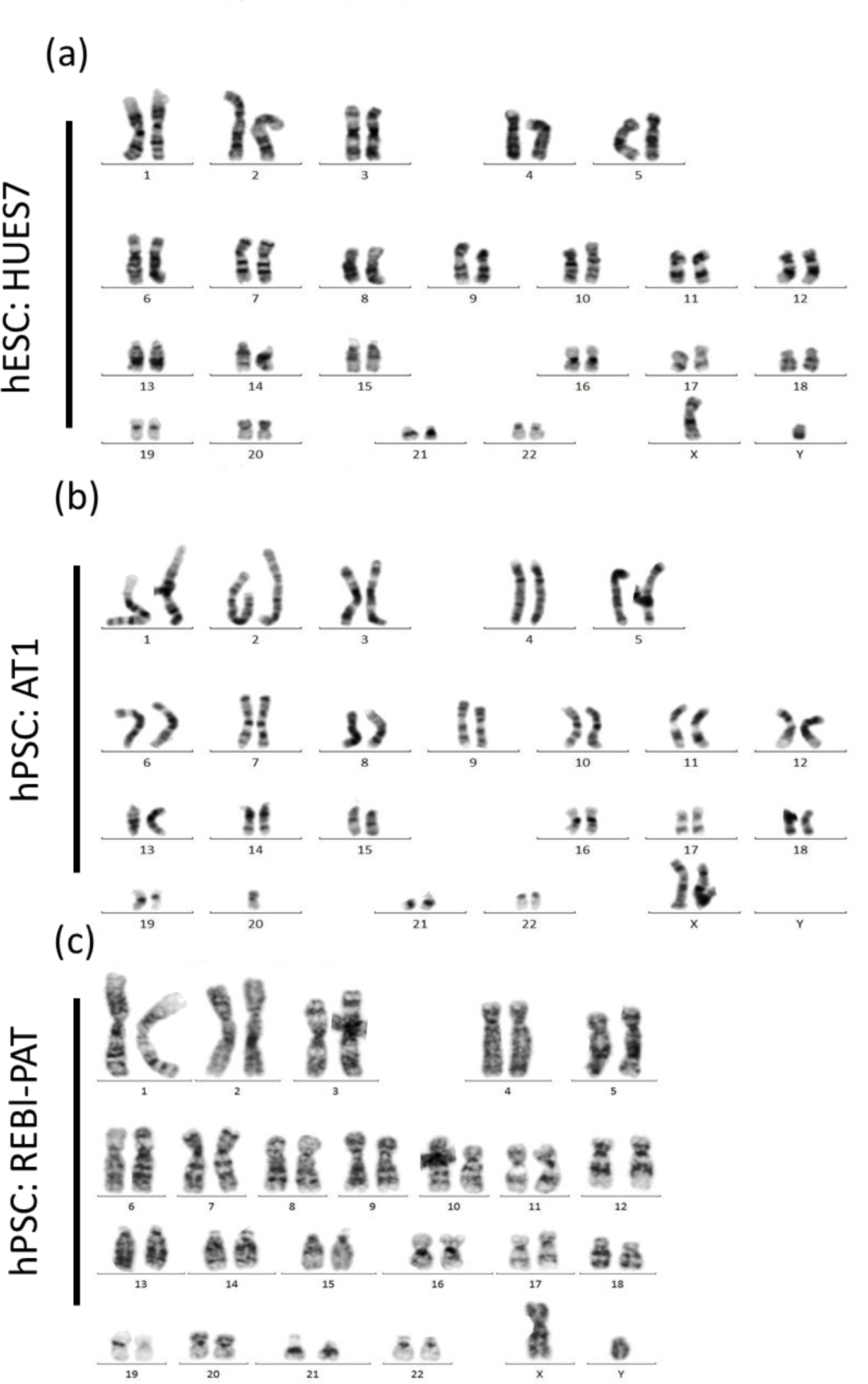
Karyograms observed after 5 serial passages on poly (TCDMDA-blend-BA) for (a) hESC HUES7 (46,XY), (b) hiPSC AT1 (46, XX) and (c) hiPSC REBl-PAT (46, XY) cultured in E8 medium.

**Figure S9:**
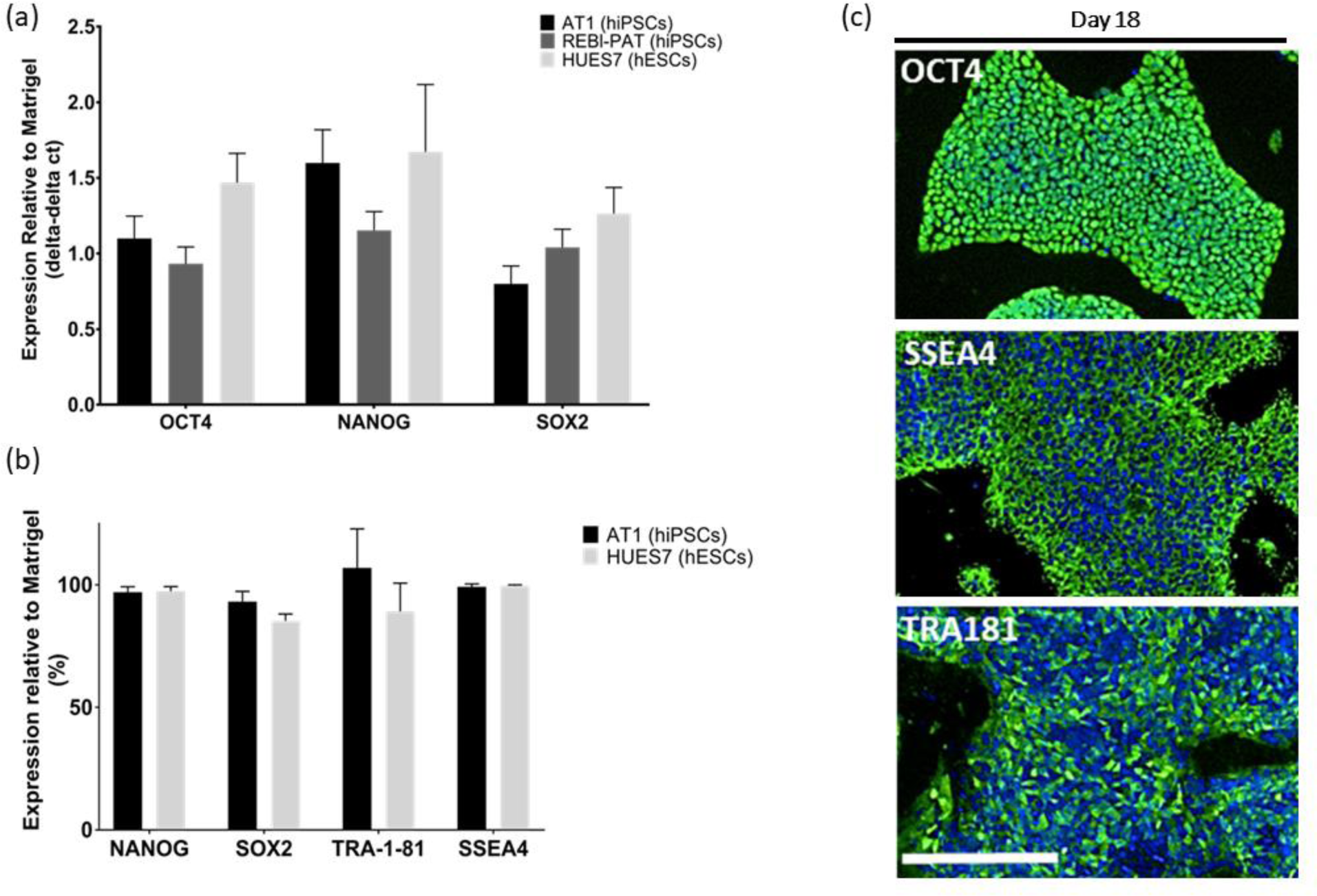
hPSCs (hiPSC AT1 and REBl-PAT lines and hESC HUES7 line) were assessed for pluripotency markers after 18 days (5 serial passages) on poly(TCDMDA-blend-BA) by (a) quantitative real-time PCR (AT1, REBl-Pat and HUES7 hPSCs), (b) flow cytometry (AT1 and HUES7 hPSCs) (c) and immunostaining ReBl-PAT) for pluripotent markers (including NANOG, SOX2, OCT4, SSEA4 and TRA181). Scale bar represents 200μm.

**Figure S10:**
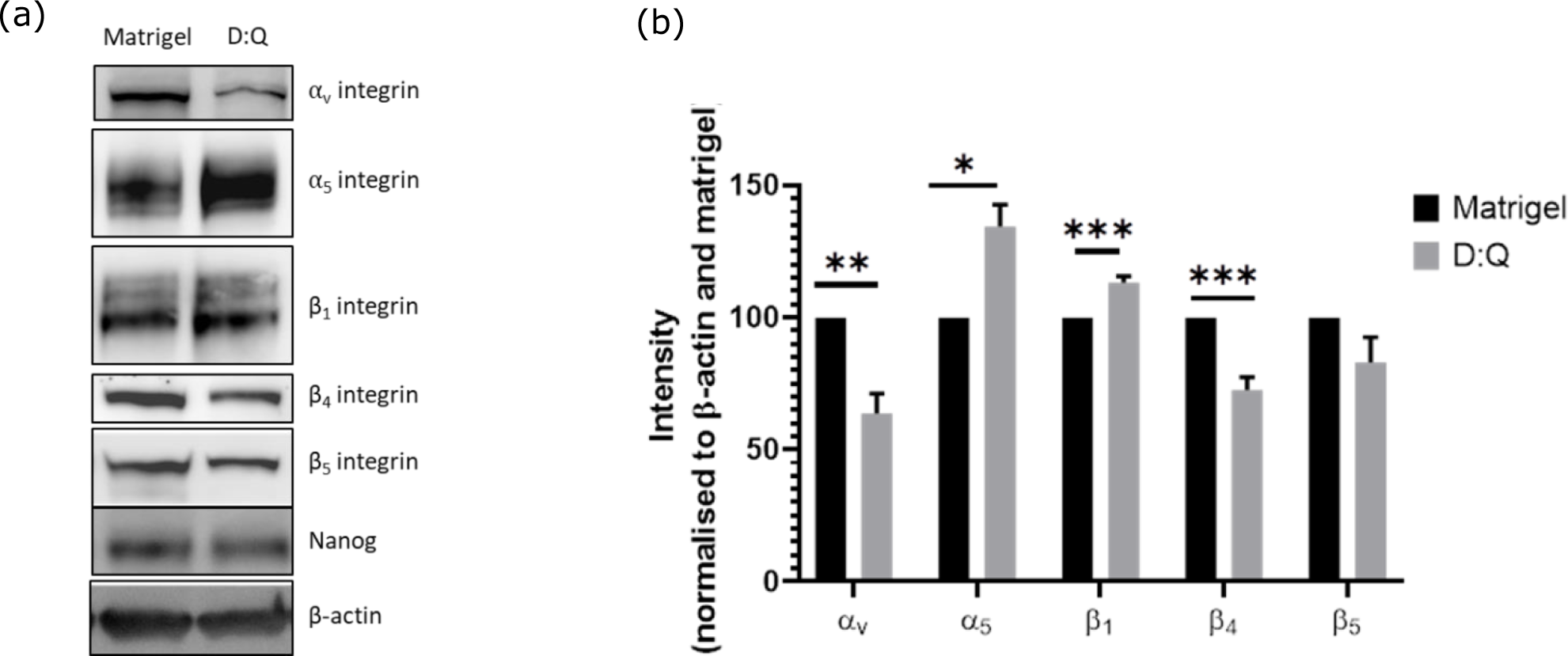
Protein expression of integrin subunits in hiPSC AT1 cells cultured on Matrigel and D:Q (poly(TCDMDA-blend-BA) for at least three serial passages assessed by western blot analysis. (a) Representative images of Western Blotting bands for integrin subunits ⍺v, ⍺5, β1, β4, β5; stem cell marker Nanog, and house-keeping protein β-actin, (n=3). (b) Quantification of band intensity for integrin expression in AT-1 hiPSCs (n⩾3), bars show Mean ± STDEV; black bars show Matrigel control and grey bars show AT-1 on the hit Polymer. Unpaired t-test were performed, and statistical significance is represented as: *P<0.033, **P<0.002, ***P<0.001.

**Figure S11:**
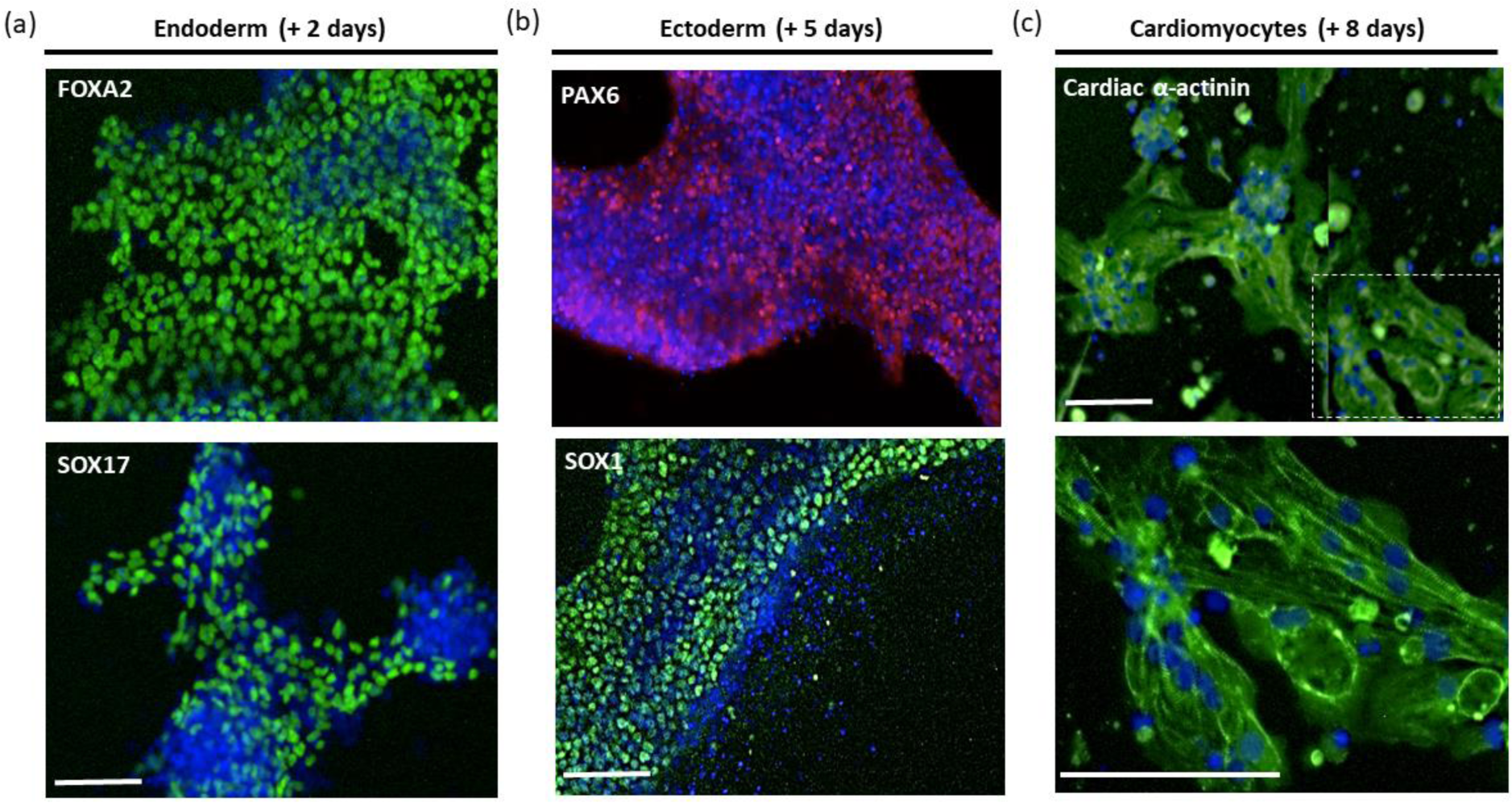
Tri-lineage differentiation of REBl-PAT hPSCs cultured on poly (TCDMDA-blend-BA) for five passages. (a) Definitive endoderm differentiation induced early-stage marker expression of FOXA2 and SOX17 after 2 days. (b) Ectoderm differentiation induced neurogenesis marker expression after 5 days. (c) Mesoderm differentiation induced positive α-actinin expression after 8 days. Scale bars represent 100μm.

**Table S1:** Full list of monomers included for microarray screens with acronyms and full IUPAC names.

| Acronym | Name |
| --- | --- |
| 13BDDA | Butanediol-1,3 diacrylate |
| THFuA | Tetrahydrofurfuryl acrylate |
| EGDPEA | Ethylene glycol dicyclopentenyl ether acrylate |
| MAAH | Methacrylic anhydride |
| MAPtMA | Methacrylamidopropyltrimethylammonium chloride, |
| MAEA | Methacryloyloxyethyl acetoacetate |
| 13BDDMA | 1,3-Butanediol dimethacrylate |
| EGDA | Ethylene glycol diacrylate |
| TMPETA | Trimethylolpropane ethoxylate triacrylate |
| TMCHMA | Trimethylcyclohexyl methacrylate |
| TMOBDA | Trimethylolpropane benzoate diacrylate |
| DMAEMA | Dimethylamino-ethyl methacrylate |
| BDDA | Butanediol diacrylate |
| SolA | Solketal acrylate |
| HBOPBA | Hexanediylbis[oxy(2-hydroxy-3,1-propanediyl)] bisacrylate |
| E3GDA | Triethylene glycol diacrylate |
| DVAd | Divinyl Adipate |
| PDDMA | 1,5-Pentanediol dimethacrylate |
| TAHTA | 1,3,5-Triacryloylhexahydro-1,3,5-triazine |
| CNEA | Cyanoethyl acrylate |
| EGDCMA | Ethylene glycol dicyclopentenyl ether methacrylate |
| OFPA | Octafluoropentyl acrylate |
| DPEPHA | Dipentaerythritol penta/hexa-acrylate |
| ZrBNCTA | Zirconium bromonorbornanelactone carboxylate triacrylate |
| HEODA | Hexanediol ethoxylate diacrylate |
| HDMPDA | Hydroxy-2,2-dimethylpropyl 3-hydroxy-2,2-dimethylpropionate diacrylate |
| PETrA | Pentaerythritol triacrylate |
| MAEACl | [2-(Methacryloyloxyethyl)trimethylammonium chloride solution |
| DEGDA | Di(ethylene glycol) diacrylate |
| NpMA | Naphthyl methacrylate |
| TBNpMA | Tribromoneopentyl methacrylate |
| 14BDDMA | 1,4-Butanediol dimethacrylate |
| TMHA | Trimethylhexyl acrylate |
| mMAOEM | mono-2-(Methacryloyloxy)ethyl maleate |
| DMAPA | Dimethylamino-propyl acrylate |
| DDDMA | 1,10-Decanediol dimethacrylate |
| NGDA | Neopentyl glycol diacrylate |
| TAIC | Tris[2-(acryloyloxy)ethyl] isocyanurate |
| tBAEMA | Tert-butylamino-ethyl methacrylate |
| NpA | Naphthyl acrylate |
| EGPEA | Ethylene glycol phenyl ether acrylate |
| AEMA.C | 2-Aminoethyl methacrylate hydrochloride, |
| LaA | Lauryl acrylate |
| BAPODA | Bisphenol A propoxylate diacrylate |
| APMAm.C | N-(3-Aminopropyl)methacrylamide hydrochloride |
| HFPDA | Hexafluoropent-1,5-diyl diacrylate |
| tBCHA | Tert-butylcyclohexylacrylate |
| TMPTA | Trimethylolpropane triacrylate |
| DFHA | Dodecafluoroheptyl acrylate |
| AOHPMA | Acryloyloxy-2-hydroxypropyl methacrylate |
| EGDMA | Ethylene glycol dimethacrylate |
| NDDMA | 1,9-Nonanediol dimethacrylate |
| PhA | Phenyl acrylate |
| TPGDA | Tri(propylene glycol) diacrylate |
| BnA | Benzyl acrylate |
| HDDMA | 1,6-Hexanediol dimethacrylate, |
| FuMA | Furfuryl methacrylate |
| BzHPEA | Benzoyl-3-hydroxy-phenoxy)ethyl acrylate |
| ExA | Epoxidized acrylate |
| CHMA | Cyclohexyl methacrylate |
| TMOPTMA | 1,1,1-Trimethylolpropane trimethacrylate |
| BPAPGDA | Bisphenol A propoxylate glycerolate diacrylate |
| iBMA/tBAEMA | Isobornyl methacrylate:Tert-butylamino-ethyl methacrylate 2:1 |
| PhMA | Phenyl methacrylate |
| BHMA | Benzhydryl methacrylate |
| pEGDMA | Poly(ethylene glycol) (600) dimethacrylate |
| DEGEEA | Di(ethylene glycol) ethyl ether acrylate |
| BAGDA | Bisphenol A glycerolate diacrylate |
| ODA | Octadecyl acrylate |
| iCEMA | Isocyanatoethyl methacrylate |
| BHMOPhP | 2,2-Bis[4-(2-hydroxy-3-methacryloxypropoxy)phenyl]propane |
| DMPMAm | N-[3-(Dimethylamino)propyl]methacrylamide |
| GPOTA | Glycerol propoxylate triacrylate |
| HFiPA | Hexafluoroisopropyl acrylate |
| iBMA | Isobornyl methacrylate |
| BnMA | Benzyl methacrylate |
| HPhOPA | Hydroxy-3-phenoxypropyl acrylate |
| iOA | Isooctyl acrylate |
| PDA | 1,4-Phenylene diacrylate |
| PETA | Pentaerythritol tetraacrylate |
| TEGDA | Tetra(ethylene glycol) diacrylate |
| GDMA | Glycerol dimethacrylate |
| HDMA | 1-Hexadecyl methacrylate |
| TCMDMA | Tricyclodecane-dimethanol diacrylate |
| MAHBP | 4-Methacryloxy-2-hydroxybenzophenone |
| BTHPhMA | Benzotriazol-2-yl)-4-hydroxyphenyl]ethyl methacrylate |
| DMA | Decyl methacrylate |
| NGPDA | Neopentyl glycol propoxylate diacrylate |
| DMEMAm | N-[2-(N,N-Dimethylamino)ethyl]methacrylamide |
| DEAEA | Diethylamino ethyl acrylate |
| pEGPhEA | Poly(ethylene glycol) phenyl ether acrylate |
| PhEMA | 2-Phenylethyl methacrylate |
| pPGDMA | Poly(propylene glycol) (400) dimethacrylate |
| MAAHS | Methacrylic acid N-hydroxysuccinimide ester |
| PPDDA | 3-phenoxypropane-1,2-diyl diacrylate |
| PHPhMA | 3-Phenoxy 2 hydroxy propyl methacrylate |
| HEODA/EEMA | Hexanediol ethoxylate diacrylate:Ethoxyethyl methacrylate 2:1 |
| HPHPBAH | Hydroxypivalyl hydroxypivalate bis[6-(acryloyloxy)hexanoate] |
| PFPPhA | Pentafluorophenyl acrylate |
| DEAEMA | Diethylaminoethyl methacrylate |
| TBPhA | 2,4,6-Tribromophenyl acrylate |
| PMMA | 1-Pyrenylmethyl methacrylate |
| MAPU | 2-methacryloxyethyl phenyl urethane |
| NBMA | Norbornyl methacrylate |
| PhMAm | N-Phenylmethacrylamide |
| DEGEHA | Di(ethylene glycol) 2-ethylhexyl ether acrylate |
| HBMA | Hydroxybutyl methacrylate |
| pPGMEA | Poly(propylene glycol) methyl ether acrylate |
| iDMA | Isodecyl methacrylate |
| DiPEMA | 2-Diisopropylaminoethyl methacrylate |
| AEMAm.C | N-(2-aminoethyl) methacrylamide hydrochloride |
| HPMAP | Hydroxypropyl 2-(methacryloyloxy)ethyl phthalate |
| BPDMA | Bisphenol A dimethacrylate |
| MTEMA | Methylthioethyl methacrylate |
| PEDAM | Pentaerythritol diacrylate monostearate |
| MHMB | Methyl 3-hydroxy-2-methylenebutyrate |
| EG3DMA | Tri(ethylene glycol) dimethacrylate |
| HDFHUA | Heptadecafluoro-2-hydroxyundecyl acrylate |
| HPA | Hydroxypropyl acrylate |
| NaPhA | Sodium 3-phenyl-acrylate |
| ZrCEA | Zirconium carboxyethyl acrylate |
| BPEODA | Bisphenol A ethoxylate diacrylate |
| COEA | 2-Cinnamoyloxyethyl acrylate |
| DEGDMA | Diethylene glycol dimethacrylate |
| OFHMA | Octafluoro-2-hydroxy-6-(trifluoromethyl)heptyl methacrylate |
| iBuMA | Isobutyl methacrylate |
| GMA | Glycidyl methacrylate |
| iDA | Isodecyl acrylate |
| SPAK | Sulfopropyl acrylate potassium salt |
| BFEODA | Bisphenol F ethoxylate diacrylate |
| BnPA | Benzyl 2-n-propyl acrylate |
| CzEA | Carbazol-9-yl ethyl acrylate |
| tBCHMA | Tertbutylcyclohexyl methacrylate |
| TFPMA | Tetrafluoropropyl methacrylate |
| MA | Methyl acrylate |
| TDFOcA | Tridecafluorooctyl acrylate |
| MAETA | 4-Methacryloxyethyl trimellitic anhydride |
| DVSeb | Divinyl sebacate |
| TMPOTA | Trimethylolpropane propoxylate triacrylate |
| BMENBC | Bis(2-methacryloxyethyl) N,N'-1,9-nonylene biscarbamate |
| NBnMA | o-Nitrobenzyl methacrylate |
| nOcMA | n-Octyl methacrylate, |
| SMA | Stearyl methacrylate |
| HFHUMA | Hexadecafluoro-2-hydroxy-10-(trifluoromethyl)undecyl methacrylate |
| MAEP | Monoacryloxyethyl phosphate |
| CHA | Cyclohexyl acrylate |
| iBOA | Isobornyl acrylate |
| THFuMA | Tetrahydrofurfuryl methacrylate |
| DMAEA | Dimethylamino-ethyl acrylate |
| PhEA | 2-Phenylethyl acrylate |
| PAHEMA | Phosphoric acid 2-hydroxyethyl methacrylate ester |
| BOEMA | Butoxyethyl methacrylate |
| HDFDA | Heptadecafluorodecyl acrylate |
| HFIPMA | Hexafluoroisopropyl methacrylate |
| BMA | Butyl methacrylate |
| DMPAm | N-[3-(Dimethylamino)propyl]acrylamide |
| GDGDA | Glycerol 1,3-diglycerolate diacrylate |
| EHMA | Ethylhexyl methacrylate |
| DFFMOA | Dodecafluoro-7-(trifluoromethyl)-octyl acrylate |
| BAC | N,N'-Bis(acryloyl)cystamine |
| HEAm | N-Hydroxyethyl acrylamide |
| mMAOES | mono-2-(Methacryloyloxy)ethyl succinate |
| BA | Butyl acrylate |
| BMAM | N-Benzylmethacrylamide |
| FMHPNMA | Trifluoro-2'-(trifluoromethyl)-2'-hydroxypropyl]-3-norbornyl methacrylate |
| MMA | Methyl methacrylate |
| CHPMA | Chloro-2-hydroxy-propyl methacrylate |
| pPGDA | Poly(propylene glycol) diacrylate |
| BACOEa | Butylamino carbonyl oxy ethyl acrylate |
| EOEA | Ethoxyethyl acrylate |
| iBA | Isobutyl acrylate |
| SPMAK | 3-Sulfopropyl methacrylate potassium salt |
| DHPA | 2,3-dihydroxypropyl acrylate |
| F6BMA | Hexafluorobutyl methacrylate |
| pFDA | Perfluorodecyl acrylate |
| IBESMA | 1,7,7-trimethylbicyclo[2.2.1]heptan-2-yl 6-(methacryloyloxy)-4-oxohexanoate |
| HDFDMA | Heptadecafluorodecyl methacrylate |
| TFCAm | 7-[4-(Trifluoromethyl)coumarin]acrylamide |
| AODMBA | (R)-α-Acryloyloxy-$\beta,\beta$-dimethyl-γ-butyrolactone |
| PA | Propargyl acrylate |
| OFPMA | Octafluoropentyl methacrylate |
| iBOMAm | N-(Isobutoxymethyl)acrylamide |
| BAPA | 1,4-Bis(acryloyl)piperazine |
| DFHNMA | Dodecafluoro-2-hydroxy-8-(trifluoromethyl)nonyl methacrylate |
| F6BA | Hexafluorobutyl acrylate |
| MAEPC | 2-Methacryloyloxyethyl phosphorylcholine |
| pPGNEA | Poly(propylene glycol) 4-nonylphenyl ether acrylate |
| SEMA | 2-Sulfoethyl methacrylate |
| VMA | Vinyl methacrylate |
| HMA | Hexyl methacrylate |
| EbCNA | Ethyl-cis-B-cyano-acrylate |
| THMMAm | N-[Tris(hydroxymethyl)methyl]acrylamide |
| HA | Hexyl acrylate |
| tBMAM | N-tert-Butylmethacrylamide |
| HTFDA | Hexadecafluoro-9-(trifluoromethyl)decyl acrylate |
| AMA | Allyl methacrylate |
| EEMA | Ethoxyethyl methacrylate |
| EHA | Ethylhexyl acrylate |
| FuMA/tBAEMA | FuMA/tBAEMA |
| PMAM | N-(Phthalimidomethyl)acrylamide |
| tBMA | Tert-butyl methacrylate |
| TMBAm | N-(1,1,3,3-Tetramethylbutyl)acrylamide |
| DEGMA | Di(ethylene glycol) methyl ether methacrylate |
| TBPMA | Tribromophenyl methacrylate |
| EGMMA | Ethylene glycol methyl ether methacrylate |
| EEA | Ethyl 2-ethylacrylate |
| LMA | Lauryl methacrylate |
| MPDSAHA | Methacryloylamino)propyl]dimethyl(3-sulfopropyl)ammonium hydroxide inner salt |
| AnMA | Anthracenylmethylacrylate |
| EBAM | N,N'-Ethylenebisacrylamide |
| F7BA | Heptafluorobutyl acrylate |
| HFDA | Heneicosfluorododecyl acrylate |
| HPMAm | N-(2-Hydroxypropyl)methacrylamide |
| MEDMSAH | [2-(Methacryloyloxy)ethyl]dimethyl-(3-sulfopropyl) ammonium hydroxide |
| PBPhMA | Pentabromophenyl methacrylate |
| PFPPhMA | Pentafluorophenyl methacrylate |
| HBA | Hydroxybutyl acrylate |
| PPPDMA | PEO(5800)-b-PPO(3000)-b-PEO(5800) dimethacrylate |
| tBA | N-tert-Butylacrylamide |
| CEA | Carboxyethyl acrylate |
| HfCEA | Hafnium carboxyethyl acrylate |
| TMPDAE | Trimethyl propane diallyl ether |
| TPhMAm | N-(Triphenylmethyl)methacrylamide |
| DAAM | N,N-Diallylacrylamide |
| EMA | Ethyl methacrylate |
| EPA | Ethyl 2-propylacrylate |
| HMBMAm | N,N'-Hexamethylenebis(methacrylamide) |
| TDFOMA | Tridecafluorooctyl methacrylate |
| DRA | Disperse red 1 acrylate |
| HPhMA | N-(4-Hydroxyphenyl)methacrylamide |
| MAL | Methacryloyl-L-Lysine |
| NDMAm | N-Dodecylmethacrylamide |
| PBBA | Pentabromobenzyl acrylate |
| pEGMEMA | Poly(ethylene glycol) methyl ether methacrylate |
| DMMAm | N,N-Dimethylmethacrylamide |
| DYA | Disperse yellow 7 acrylate |
| EG4DMA | Tetraethylene glycol dimethacrylate |
| Mam | Methacrylamide |
| pPGA | Poly(propylene glycol) acrylate |
| DOAm | Disperse Orange 3 acrylamide |
| HPMA | Hydroxypropyl methacrylate |
| BOMAm | N-(Butoxymethyl)acrylamide |
| NMEMA | 2-N-Morpholinoethyl methacrylate |
| tBA | Tert-butyl acrylate |
| BMAOEP | Bis[2-(methacryloyloxy)ethyl] phosphate |
| ECNTA | Ethyl-2-cyano-3-(2-thienyl)acrylate |
| F7BMA | Heptafluorobutyl methacrylate |
| NAM | N-Acryloylmorpholine |
| pEGMEA | Poly(ethylene glycol) methyl ether acrylate |
| PFPMA | Pentafluoropropyl methacrylate |
| EGMEA | Ethylene glycol methyl ether acrylate |
| HMAm | N-(Hydroxymethyl)acrylamide |
| iPAM | N-Isopropylacrylamide |
| GMMA | Glycerol monomethacrylate |
| AAm | Acrylamide |
| MMAm | N-Methylmethacrylamide |
| PMA | Propargyl methacrylate |
| tBEMAm | N-(3,3-dimethylbutyl)methacrylamide |
| ZrA | Zirconium acrylate |
| AA | Allyl acrylate |
| CMAOE | Caprolactone 2-(methacryloyloxy)ethyl ester |
| DMAm | N,N'-Dimethylacrylamide |
| EGPhMA | Ethylene glycol phenyl ether methacrylate |
| HEA | Hydroxyethyl acrylate |
| MBMAm | N,N'-Methylenebismethacrylamide |
| NAS | N-Acryloxysuccinimide |
| tBOCAPAm | N-(t-BOC-aminopropyl)methacrylamide |
| TEGMA | Tri(ethylene glycol) methyl ether methacrylate |
| ZnA | Zinc acrylate |
| HMBAM | N,N'-Hexamethylenebisacrylamide |
| PBPhA | Pentabromophenyl acrylate |
| PFPA | Pentafluoropropyl acrylate |
| AAcAm | Diacetone acrylamide |
| AcAPAm | N-[2-(Acryloylamino)phenyl]acrylamide |
| DHEBAM | N,N'-(1,2-Dihydroxyethylene)bisacrylamide |
| EA | Ethyl acrylate |
| EaNIA | Ethyl trans-a-cyano-3-indole-acrylate |
| GA | Glycidyl acrylate |
| MAA | Methyl 2-acetamidoacrylate |
| MBAm | N,N'-Methylenebisacrylamide |
| MOPAm | N-(3-Methoxypropyl)acrylamide |
| NPhPMA | Nitrophenyl-2-pyrrolidonemethyl acrylate |
| pEGDA | Polyethylene glycol diacrylate |
| pEGMA | Poly(ethylene glycol) methacrylate |

**Table S2:** hPSC attachment on monomer screen at 24 h ranked (high to low) by total cell number (DAPI nuclei count) and OCT4+ nuclei count.

| Rank order | DAPI count |  |  | OCT4+ count |  |  |
| --- | --- | --- | --- | --- | --- | --- |
|  | Polymer ID | Mean | STDEV | Polymer ID | Mean | STDEV |
| 1 | **13BDDA** | 60 | 44 | **13BDDA** | 56 | 41 |
| 2 | **THFuA** | 55 | 35 | **THFuA** | 52 | 32 |
| 3 | **EGDPEA** | 51 | 22 | **EGDPEA** | 48 | 21 |
| 4 | **MAAH** | 49 | 20 | **MAAH** | 47 | 20 |
| 5 | **MAPtMA** | 43 | 39 | **MAPtMA** | 40 | 36 |
| 6 | **TMPETA** | 42 | 26 | **MAEA** | 38 | 25 |
| 7 | **TMOBDA** | 40 | 22 | **13BDDMA** | 38 | 53 |
| 8 | **BDDA** | 40 | 43 | **EGDA** | 37 | 68 |
| 9 | **MAEA** | 39 | 26 | **TMPETA** | 37 | 23 |
| 10 | **13BDDMA** | 39 | 55 | **TMCHMA** | 36 | 32 |
| 11 | **TMCHMA** | 38 | 34 | **TMOBDA** | 36 | 20 |
| 12 | **EGDA** | 38 | 70 | **DMAEMA** | 36 | 37 |
| 13 | **DMAEMA** | 37 | 37 | **BDDA** | 34 | 38 |
| 14 | **DVAd** | 35 | 18 | **SoIA** | 32 | 31 |
| 15 | **E3GDA** | 35 | 26 | **HBOPBA** | 31 | 15 |
| 16 | **SoIA** | 35 | 34 | **E3GDA** | 31 | 25 |
| 17 | **PDDMA** | 34 | 26 | **DVAd** | 31 | 16 |
| 18 | **DMEMAm** | 33 | 24 | **PDDMA** | 30 | 24 |
| 19 | **HBOPBA** | 32 | 14 | **TAHTA** | 30 | 25 |
| 20 | **TAHTA** | 31 | 26 | **CNEA** | 29 | 22 |
| 21 | **MAEACI** | 31 | 21 | **EGDCMA** | 28 | 22 |
| 22 | **DMAPA** | 30 | 32 | **OFPA** | 28 | 25 |
| 23 | CNEA | 30 | 24 | DPEPHA | 27 | 31 |
| 24 | EGDCMA | 29 | 22 | ZrBNCTA | 26 | 16 |
| 25 | OFPA | 29 | 26 | HEODA | 26 | 28 |
| 26 | DPEPHA | 29 | 33 | HDMPDA | 26 | 11 |
| 27 | HDMPDA | 28 | 12 | PETrA | 25 | 26 |
| 28 | HEODA | 28 | 30 | MAEACI | 25 | 22 |
| 29 | PETrA | 28 | 30 | DEGDA | 24 | 15 |
| 30 | ZrBNCTA | 27 | 17 | NpMA | 24 | 13 |
| 31 | DEGDA | 27 | 17 | TBNpMA | 24 | 14 |
| 32 | NpMA | 26 | 15 | 14BDDMA | 24 | 24 |
| 33 | tBAEMA | 26 | 51 | TMHA | 24 | 43 |
| 34 | 14BDDMA | 26 | 26 | mMAOEM | 23 | 15 |
| 35 | TBNpMA | 25 | 15 | DMAPA | 23 | 30 |
| 36 | NGDA | 24 | 25 | DDDMA | 23 | 21 |
| 37 | TMHA | 24 | 45 | NGDA | 23 | 24 |
| 38 | mMAOEM | 24 | 16 | TAIC | 22 | 29 |
| 39 | DMPMAm | 23 | 9 | tBAEMA | 22 | 43 |
| 40 | NpA | 23 | 22 | NpA | 21 | 20 |
| 41 | DDDMA | 23 | 21 | EGPEA | 21 | 20 |
| 42 | TAIC | 22 | 29 | AEMA.C | 20 | 22 |
| 43 | EGPEA | 22 | 20 | LaA | 19 | 22 |
| 44 | TMPTA | 22 | 21 | BAPODA | 19 | 16 |
| 45 | AEMA.C | 22 | 24 | APMAm.C | 19 | 30 |
| 46 | HFiPA | 21 | 20 | HFPDA | 19 | 12 |
| 47 | DFHA | 20 | 12 | tBCHA | 18 | 20 |
| 48 | APMAm.C | 20 | 32 | TMPTA | 18 | 18 |
| 49 | NDDMA | 20 | 19 | DFHA | 18 | 11 |
| 50 | tBCHA | 20 | 21 | AOHPMA | 17 | 21 |
| 51 | BAPODA | 20 | 16 | EGDMA | 17 | 20 |
| 52 | LaA | 19 | 22 | NDDMA | 17 | 16 |
| 53 | HFPDA | 19 | 12 | PhA | 17 | 24 |
| 54 | BzHPEA | 19 | 19 | TPGDA | 17 | 21 |
| 55 | TPGDA | 19 | 21 | BnA | 17 | 15 |
| 56 | BnA | 19 | 17 | HDDMA | 17 | 13 |
| 57 | PhA | 18 | 25 | FuMA | 17 | 27 |
| 58 | HDDMA | 18 | 15 | BzHPEA | 17 | 16 |
| 59 | pEGDMA | 18 | 17 | ExA | 16 | 26 |
| 60 | EGDMA | 18 | 21 | CHMA | 16 | 16 |
| 61 | FuMA | 18 | 29 | TMOPTMA | 16 | 11 |
| 62 | PhMA | 18 | 21 | BPAPGDA | 16 | 26 |
| 63 | AOHPMA | 18 | 21 | iBMA/tBAEMA | 16 | 27 |
| 64 | CHMA | 17 | 16 | PhMA | 15 | 19 |
| 65 | BAGDA | 17 | 25 | BHMA | 15 | 19 |
| 66 | ExA | 16 | 26 | pEGDMA | 15 | 15 |
| 67 | BHMOPhP | 16 | 12 | DEGEEA | 15 | 22 |
| 68 | **TMOPTMA** | 16 | 12 | **BAGDA** | 15 | 22 |
| 69 | **iBMA/tBAEMA** | 16 | 27 | **ODA** | 15 | 8 |
| 70 | **ODA** | 16 | 9 | **iCEMA** | 14 | 12 |
| 71 | **BHMA** | 16 | 19 | **BHMOPhP** | 14 | 11 |
| 72 | **BPAPGDA** | 16 | 27 | **DMPMAm** | 14 | 7 |
| 73 | **GPOTA** | 16 | 9 | **GPOTA** | 14 | 8 |
| 74 | **iBMA** | 15 | 29 | **HFiPA** | 14 | 16 |
| 75 | **BnMA** | 15 | 14 | **iBMA** | 13 | 25 |
| 76 | **DEGEEA** | 15 | 22 | **BnMA** | 13 | 12 |
| 77 | **iCEMA** | 15 | 12 | **HPhOPA** | 13 | 18 |
| 78 | **NGPDA** | 15 | 18 | **iOA** | 13 | 11 |
| 79 | **DMA** | 15 | 19 | **PDA** | 13 | 9 |
| 80 | **MAHBP** | 15 | 7 | **PETA** | 13 | 7 |
| 81 | **HDMA** | 14 | 26 | **TEGDA** | 13 | 18 |
| 82 | **PETA** | 14 | 8 | **GDMA** | 13 | 9 |
| 83 | **DEAEA** | 14 | 20 | **HDMA** | 13 | 24 |
| 84 | **HPhOPA** | 14 | 20 | **TCDMDA** | 13 | 17 |
| 85 | **PHPMA** | 14 | 17 | **MAHBP** | 12 | 7 |
| 86 | **TCDMDA** | 14 | 19 | **BTHPhMA** | 12 | 11 |
| 87 | **PDA** | 14 | 10 | **DMA** | 12 | 17 |
| 88 | **GDMA** | 13 | 10 | **NGPDA** | 12 | 14 |
| 89 | **iOA** | 13 | 12 | **DMEMAm** | 12 | 11 |
| 90 | **TEGDA** | 13 | 19 | **DEAEA** | 12 | 17 |
| 91 | **MAAHS** | 13 | 23 | **pEGPhEA** | 12 | 17 |
| 92 | **HPHPBAH** | 13 | 10 | **PhEMA** | 12 | 19 |
| 93 | **BTHPhMA** | 13 | 11 | **pPGDMA** | 12 | 13 |
| 94 | **TBPhA** | 13 | 13 | **MAAHS** | 12 | 20 |
| 95 | **HEODA/EEMA** | 12 | 22 | **PPDDA** | 11 | 14 |
| 96 | **PhEMA** | 12 | 20 | **PHPMA** | 11 | 14 |
| 97 | **pEGPhEA** | 12 | 18 | **HEODA/EEMA** | 11 | 20 |
| 98 | **PPDDA** | 12 | 15 | **HPHPBAH** | 11 | 9 |
| 99 | **pPGDMA** | 12 | 13 | **PFPhA** | 11 | 9 |
| 100 | **iDMA** | 12 | 20 | **DEAEMA** | 11 | 12 |
| 101 | **HBMA** | 11 | 34 | **TBPhA** | 11 | 13 |
| 102 | **PFPhA** | 11 | 9 | **PMMA** | 11 | 15 |
| 103 | **DEGEHA** | 11 | 16 | **MAPU** | 10 | 12 |
| 104 | **MAPU** | 11 | 14 | **NBMA** | 10 | 15 |
| 105 | **NBMA** | 11 | 15 | **PhMAm** | 10 | 14 |
| 106 | **PhMAm** | 11 | 15 | **DEGEHA** | 10 | 15 |
| 107 | **PMMA** | 11 | 15 | **HBMA** | 10 | 30 |
| 108 | **DEAEMA** | 11 | 13 | **pPGMEA** | 10 | 14 |
| 109 | **pPGMEA** | 11 | 15 | **iDMA** | 10 | 17 |
| 110 | **HPA** | 10 | 17 | **DiPEMA** | 10 | 11 |
| 111 | **PEDAM** | 10 | 24 | **AEMAm.C** | 9 | 15 |
| 112 | **MTEMA** | 10 | 13 | **HPMAP** | 9 | 12 |
| 113 | **DiPEMA** | 10 | 11 | **BPDMA** | 9 | 10 |
| 114 | **AEMAm.C** | 10 | 16 | **MTEMA** | 9 | 11 |
| 115 | **HPMAP** | 10 | 12 | **PEDAM** | 8 | 19 |
| 116 | **EG3DMA** | 9 | 15 | **MHMB** | 8 | 8 |
| 117 | **BPDMA** | 9 | 10 | **EG3DMA** | 8 | 13 |
| 118 | **HDFHUA** | 9 | 15 | **HDFHUA** | 8 | 14 |
| 119 | **COEA** | 9 | 12 | **HPA** | 8 | 16 |
| 120 | **DEGDMA** | 9 | 9 | **NaPhA** | 8 | 8 |
| 121 | **DMPAm** | 9 | 12 | **ZrCEA** | 8 | 8 |
| 122 | **iDA** | 9 | 11 | **BPEODA** | 8 | 9 |
| 123 | **MHMB** | 9 | 8 | **COEA** | 8 | 11 |
| 124 | **BPEODA** | 9 | 10 | **DEGDMA** | 8 | 8 |
| 125 | **NaPhA** | 8 | 8 | **OFHMA** | 8 | 8 |
| 126 | **iBuMA** | 8 | 10 | **iBuMA** | 7 | 9 |
| 127 | **SPAK** | 8 | 6 | **GMA** | 7 | 14 |
| 128 | **ZrCEA** | 8 | 8 | **iDA** | 7 | 9 |
| 129 | **OFHMA** | 8 | 8 | **SPAK** | 7 | 6 |
| 130 | **TDFOcA** | 8 | 8 | **BFEODA** | 7 | 7 |
| 131 | **GMA** | 7 | 14 | **BnPA** | 7 | 7 |
| 132 | **MA** | 7 | 10 | **CzEA** | 7 | 7 |
| 133 | **TFPMA** | 7 | 10 | **tBCHMA** | 7 | 6 |
| 134 | **BFEODA** | 7 | 7 | **TFPMA** | 7 | 10 |
| 135 | **BnPA** | 7 | 7 | **MA** | 7 | 9 |
| 136 | **MAETA** | 7 | 9 | **TDFOcA** | 7 | 7 |
| 137 | **tBCHMA** | 7 | 7 | **MAETA** | 6 | 9 |
| 138 | **CzEA** | 7 | 7 | **DVSeb** | 6 | 6 |
| 139 | **SMA** | 7 | 10 | **TMPOTA** | 6 | 12 |
| 140 | **BMENBC** | 7 | 2 | **BMENBC** | 6 | 2 |
| 141 | **DVSeb** | 7 | 6 | **NBnMA** | 6 | 18 |
| 142 | **HFHUMA** | 7 | 5 | **nOcMA** | 6 | 9 |
| 143 | **PAHEMA** | 7 | 7 | **SMA** | 6 | 9 |
| 144 | **HDFDA** | 6 | 8 | **HFHUMA** | 6 | 5 |
| 145 | **TMPOTA** | 6 | 12 | **MAEP** | 6 | 8 |
| 146 | **nOcMA** | 6 | 9 | **CHA** | 5 | 8 |
| 147 | **iBOA** | 6 | 7 | **iBOA** | 5 | 6 |
| 148 | **NBnMA** | 6 | 18 | **THFuMA** | 5 | 7 |
| 149 | **PhEA** | 6 | 14 | **DMAEA** | 5 | 11 |
| 150 | **BOEMA** | 6 | 8 | **PhEA** | 5 | 12 |
| 151 | **BMA** | 6 | 10 | **PAHEMA** | 5 | 5 |
| 152 | **THFuMA** | 6 | 7 | **BOEMA** | 5 | 7 |
| 153 | **CHA** | 6 | 9 | **HDFDA** | 5 | 5 |
| 154 | **MAEP** | 6 | 8 | **HFiPMA** | 5 | 7 |
| 155 | **GDGDA** | 5 | 5 | **BMA** | 5 | 8 |
| 156 | **HEAm** | 5 | 7 | **DMPAm** | 5 | 6 |
| 157 | **DMAEA** | 5 | 11 | **GDGDA** | 5 | 4 |
| 158 | **BMAM** | 5 | 6 | **EHMA** | 4 | 4 |
| 159 | **HFIPMA** | 5 | 7 | **DFFMOA** | 4 | 7 |
| 160 | **DFFMOA** | 5 | 8 | **BAC** | 4 | 5 |
| 161 | **BACOEa** | 5 | 6 | **HEAm** | 4 | 6 |
| 162 | **EHMA** | 4 | 4 | **mMAOES** | 4 | 4 |
| 163 | **BAC** | 4 | 5 | **BA** | 4 | 8 |
| 164 | **mMAOES** | 4 | 4 | **BMAM** | 4 | 6 |
| 165 | **pPGDA** | 4 | 5 | **FMHPNMA** | 4 | 4 |
| 166 | **BA** | 4 | 8 | **MMA** | 4 | 6 |
| 167 | **MMA** | 4 | 6 | **CHPMA** | 4 | 7 |
| 168 | **FMHPNMA** | 4 | 4 | **pPGDA** | 4 | 4 |
| 169 | **SPMAK** | 4 | 7 | **BACOEa** | 4 | 5 |
| 170 | **CHPMA** | 4 | 7 | **EOEA** | 4 | 6 |
| 171 | **DHPA** | 4 | 6 | **iBA** | 4 | 3 |
| 172 | **PMAM** | 4 | 6 | **SPMAK** | 3 | 6 |
| 173 | **OFPMa** | 4 | 7 | **DHPA** | 3 | 6 |
| 174 | **EOEA** | 4 | 6 | **F6BMA** | 3 | 3 |
| 175 | **iBA** | 4 | 3 | **pFDA** | 3 | 3 |
| 176 | **pFDA** | 4 | 3 | **IBESMA** | 3 | 4 |
| 177 | **TFCAM** | 4 | 5 | **HDFDMA** | 3 | 4 |
| 178 | **F6BMA** | 3 | 3 | **TFCAM** | 3 | 5 |
| 179 | **IBESMA** | 3 | 4 | **AODMBA** | 3 | 4 |
| 180 | **HDFDMA** | 3 | 4 | **PA** | 3 | 6 |
| 181 | **PA** | 3 | 7 | **OFPMa** | 3 | 5 |
| 182 | **EbcNA** | 3 | 7 | **iBOMAM** | 3 | 2 |
| 183 | **AODMBA** | 3 | 4 | **BAPA** | 3 | 3 |
| 184 | **pPGNEA** | 3 | 6 | **DFHNMA** | 3 | 3 |
| 185 | **iBOMAM** | 3 | 2 | **F6BA** | 3 | 4 |
| 186 | **HMA** | 3 | 6 | **MAEPC** | 3 | 5 |
| 187 | **SEMA** | 3 | 5 | **pPGNEA** | 3 | 6 |
| 188 | **BAPA** | 3 | 3 | **SEMA** | 3 | 5 |
| 189 | **DFHNMA** | 3 | 3 | **VMA** | 3 | 2 |
| 190 | **F6BA** | 3 | 4 | **HMA** | 2 | 5 |
| 191 | **MAEPC** | 3 | 5 | **EbcNA** | 2 | 6 |
| 192 | **EEMA** | 3 | 5 | **THMMAM** | 2 | 4 |
| 193 | **tBMAM** | 3 | 3 | **HA** | 2 | 3 |
| 194 | **VMA** | 3 | 2 | **tBMAM** | 2 | 3 |
| 195 | **HA** | 2 | 3 | **HTFDA** | 2 | 3 |
| 196 | **EHA** | 2 | 6 | **AMA** | 2 | 4 |
| 197 | **THMMAM** | 2 | 4 | **EEMA** | 2 | 5 |
| 198 | **AMA** | 2 | 4 | **EHA** | 2 | 5 |
| 199 | **EGMMA** | 2 | 4 | **FuMA/tBAEMA** | 2 | 2 |
| 200 | **HTFDA** | 2 | 3 | **PMAM** | 2 | 3 |
| 201 | **DEGMA** | 2 | 3 | **tBMA** | 2 | 4 |
| 202 | **FuMA/tBAEMA** | 2 | 2 | **TMBAM** | 2 | 4 |
| 203 | tBMA | 2 | 4 | DEGMA | 2 | 3 |
| 204 | TMBAm | 2 | 4 | TBPMA | 2 | 3 |
| 205 | TBPMA | 2 | 3 | EGMMA | 2 | 3 |
| 206 | HFDA | 2 | 3 | EEA | 2 | 2 |
| 207 | LMA | 2 | 2 | LMA | 2 | 2 |
| 208 | tBAm | 2 | 4 | MPDSAH | 2 | 2 |
| 209 | AnMA | 2 | 2 | AnMA | 2 | 2 |
| 210 | EEA | 2 | 2 | EBAM | 1 | 2 |
| 211 | MPDSAH | 2 | 2 | F7BA | 1 | 2 |
| 212 | EBAM | 2 | 2 | HFDA | 1 | 3 |
| 213 | PFPPhMA | 2 | 3 | HPMAm | 1 | 3 |
| 214 | F7BA | 1 | 2 | MEDMSAH | 1 | 2 |
| 215 | HPMAm | 1 | 3 | PBPhMA | 1 | 2 |
| 216 | MEDMSAH | 1 | 2 | PFPPhMA | 1 | 3 |
| 217 | PBPhMA | 1 | 2 | HBA | 1 | 3 |
| 218 | TMPDAE | 1 | 4 | PPPDMA | 1 | 2 |
| 219 | DAAM | 1 | 2 | tBAm | 1 | 3 |
| 220 | HBA | 1 | 3 | CEA | 1 | 3 |
| 221 | MAL | 1 | 2 | HfCEA | 1 | 3 |
| 222 | NDMAm | 1 | 2 | TMPDAE | 1 | 4 |
| 223 | PPPDMA | 1 | 2 | TPhMAm | 1 | 2 |
| 224 | CEA | 1 | 3 | DAAM | 1 | 1 |
| 225 | HfCEA | 1 | 3 | EMA | 1 | 3 |
| 226 | PBBA | 1 | 4 | EPA | 1 | 2 |
| 227 | TPhMAm | 1 | 2 | HMBMAm | 1 | 3 |
| 228 | EMA | 1 | 3 | TDFOMA | 1 | 2 |
| 229 | EPA | 1 | 2 | DRA | 1 | 2 |
| 230 | DYA | 1 | 3 | HPhMA | 1 | 2 |
| 231 | HMBMAm | 1 | 3 | MAL | 1 | 2 |
| 232 | pEGMEMA | 1 | 2 | NDMAm | 1 | 1 |
| 233 | TDFOMA | 1 | 2 | PBBA | 1 | 3 |
| 234 | DRA | 1 | 2 | pEGMEMA | 1 | 2 |
| 235 | HPhMA | 1 | 2 | DMMAm | 1 | 2 |
| 236 | pPGA | 1 | 1 | DYA | 1 | 2 |
| 237 | DHEBAM | 1 | 3 | EG4DMA | 1 | 1 |
| 238 | DMMAm | 1 | 2 | Mam | 1 | 2 |
| 239 | EG4DMA | 1 | 1 | pPGA | 1 | 1 |
| 240 | Mam | 1 | 2 | DOAm | 1 | 1 |
| 241 | DOAm | 1 | 1 | HPMA | 1 | 1 |
| 242 | HPMA | 1 | 1 | BOMAm | 1 | 2 |
| 243 | BOMAm | 1 | 2 | NMEMA | 1 | 1 |
| 244 | NMEMA | 1 | 1 | tBA | 1 | 1 |
| 245 | tBA | 1 | 1 | BMAOEP | 1 | 2 |
| 246 | BMAOEP | 1 | 2 | ECNTA | 1 | 1 |
| 247 | ECNTA | 1 | 1 | F7BMA | 1 | 2 |
| 248 | EGMEA | 1 | 1 | NAM | 1 | 2 |
| 249 | F7BMA | 1 | 2 | pEGMEA | 1 | 1 |
| 250 | NAM | 1 | 2 | PFPMA | 1 | 1 |
| 251 | pEGMEA | 1 | 1 | EGMEA | 1 | 1 |
| 252 | PFPMA | 1 | 1 | HMAm | 1 | 2 |
| 253 | ZnA | 1 | 1 | iPAM | 1 | 1 |
| 254 | HMAm | 1 | 2 | GMMA | 1 | 1 |
| 255 | iPAM | 1 | 1 | AAm | 0 | 1 |
| 256 | ZrA | 1 | 1 | MMAm | 0 | 1 |
| 257 | GMMA | 1 | 1 | PMA | 0 | 1 |
| 258 | AAm | 0 | 1 | tBEMAm | 0 | 1 |
| 259 | MMAm | 0 | 1 | ZrA | 0 | 1 |
| 260 | PMA | 0 | 1 | AA | 0 | 1 |
| 261 | tBEMAm | 0 | 1 | CMAOE | 0 | 1 |
| 262 | AA | 0 | 1 | DMAm | 0 | 1 |
| 263 | CMAOE | 0 | 1 | EGPhMA | 0 | 1 |
| 264 | DMAm | 0 | 1 | HEA | 0 | 1 |
| 265 | EGPhMA | 0 | 1 | MBMAm | 0 | 1 |
| 266 | HEA | 0 | 1 | NAS | 0 | 1 |
| 267 | MBMAm | 0 | 1 | tBOCAPAm | 0 | 1 |
| 268 | NAS | 0 | 1 | TEGMA | 0 | 1 |
| 269 | tBOCAPAm | 0 | 1 | ZnA | 0 | 1 |
| 270 | TEGMA | 0 | 1 | HMBAM | 0 | 1 |
| 271 | HMBAM | 0 | 1 | PBPhA | 0 | 1 |
| 272 | PBPhA | 0 | 1 | PFPA | 0 | 1 |
| 273 | PFPA | 0 | 1 | AAcAm | 0 | 0 |
| 274 | pEGDA | 0 | 0 | AcAPAm | 0 | 0 |
| 275 | AAcAm | 0 | 0 | DHEBAM | 0 | 0 |
| 276 | AcAPAm | 0 | 0 | EA | 0 | 0 |
| 277 | EA | 0 | 0 | EaNIA | 0 | 0 |
| 278 | EaNIA | 0 | 0 | GA | 0 | 0 |
| 279 | GA | 0 | 0 | MAA | 0 | 0 |
| 280 | MAA | 0 | 0 | MBAm | 0 | 0 |
| 281 | MBAm | 0 | 0 | MOPAm | 0 | 0 |
| 282 | MOPAm | 0 | 0 | NPhPMA | 0 | 0 |
| 283 | NPhPMA | 0 | 0 | pEGDA | 0 | 0 |
| 284 | pEGMA | 0 | 0 | pEGMA | 0 | 0 |

**Table S3.** hPSC attachment on co-polymer arrays after 24 h ranked (high-low) by OCT4+ nuclei count

| Rank order* | Letter ID | Polymer ID | Average OCT4 count | STDEV |
| --- | --- | --- | --- | --- |
| 1 | D:O | TCMDMA:MAETA | 164 | 80 |
| 2 | P:E | THFuA:EG4DMA | 153 | 153 |
| 3 | F:G | BDDA:EGDA | 151 | 90 |
| 4 | H:Q | GDMA:BA | 135 | 119 |
| 5 | P:B | THFuA:NGDA | 134 | 127 |
| 6 | P:C | THFuA:BHMOPhP | 131 | 122 |
| 7 | P:H | THFuA:GDMA | 129 | 108 |
| 8 | O:D | MAETA:TCMDMA | 124 | 96 |
| 9 | G:F | EGDA:BDDA | 123 | 85 |
| 10 | H:F | GDMA:BDDA | 123 | 87 |
| 11 | B:J | NGDA:mMAOES | 122 | 70 |
| 12 | D:G | TCMDMA:EGDA | 120 | 74 |
| 13 | F:P | BDDA:THFuA | 117 | 96 |
| 14 | H:D | GDMA:TCMDMA | 117 | 94 |
| 15 | M:C | EGDPEA:BHMOPhP | 115 | 87 |
| 16 | P | THFuA | 112 | 154 |
| 17 | C | BHMOPhP | 112 | 141 |
| 18 | D:N | TCMDMA:FuMA | 112 | 103 |
| 19 | I:B | DEAMEA:NGDA | 111 | 90 |
| 20 | L:D | HEMA:TCMDMA | 110 | 91 |
| 21 | D:P | TCMDMA:THFuA | 110 | 57 |
| 22 | L:E | HEMA:EG4DMA | 109 | 145 |
| 23 | E:A | EG4DMA:HBOPBA | 104 | 71 |
| 24 | D:Q | TCMDMA:BA | 104 | 102 |
| 25 | H:R | GDMA:TDFOMA | 103 | 106 |
| 26 | D | TCMDMA | 102 | 85 |
| 27 | D:B | TCMDMA:NGDA | 102 | 125 |
| 28 | F:J | BDDA:mMAOES | 101 | 54 |
| 29 | M:E | EGDPEA:EG4DMA | 101 | 101 |
| 30 | P:N | THFuA:FuMA | 100 | 89 |
| 31 | D:J | TCMDMA:mMAOES | 98 | 46 |
| 32 | H:C | GDMA:BHMOPhP | 98 | 64 |
| 33 | H:N | GDMA:FuMA | 98 | 41 |
| 34 | F:O | BDDA:MAETA | 98 | 51 |
| 35 | O:B | MAETA:NGDA | 98 | 127 |
| 36 | G | EGDA | 97 | 86 |
| 37 | F:K | BDDA:tBAEMA | 95 | 141 |
| 38 | L:B | HEMA:NGDA | 95 | 145 |
| 39 | B:F | NGDA:BDDA | 94 | 83 |
| 40 | H:B | GDMA:NGDA | 94 | 74 |
| 41 | B:H | NGDA:GDMA | 94 | 68 |
| 42 | B:L | NGDA:HEMA | 94 | 52 |
| 43 | O:A | MAETA:HBOPBA | 93 | 132 |
| 44 | N:F | FuMA:BDDA | 93 | 93 |
| 45 | R:G | TDFOMA:EGDA | 92 | 192 |
| 46 | P:F | THFuA:BDDA | 90 | 88 |
| 47 | H:G | GDMA:EGDA | 90 | 50 |
| 48 | O:H | MAETA:GDMA | 90 | 83 |
| 49 | I:E | DEAMEA:EG4DMA | 90 | 79 |
| 50 | N | FuMA | 90 | 96 |
| 51 | F:E | BDDA:EG4DMA | 88 | 66 |
| 52 | Q:E | BA:EG4DMA | 87 | 132 |
| 53 | D:C | TCDMDA:BHMOPhP | 87 | 74 |
| 54 | N:D | FuMA:TCDMDA | 86 | 85 |
| 55 | A:B | HBOPBA:NGDA | 86 | 107 |
| 56 | P:D | THFuA:TCDMDA | 85 | 65 |
| 57 | H:K | GDMA:tBAEMA | 85 | 57 |
| 58 | M:B | EGDPEA:NGDA | 84 | 73 |
| 59 | G:C | EGDA:BHMOPhP | 83 | 109 |
| 60 | N:I | FuMA:DEAEMA | 82 | 82 |
| 61 | P:R | THFuA:TDFOMA | 82 | 67 |
| 62 | N:C | FuMA:BHMOPhP | 82 | 72 |
| 63 | F:H | BDDA:GDMA | 81 | 68 |
| 64 | G:D | EGDA:TCDMDA | 80 | 80 |
| 65 | R:B | TDFOMA:NGDA | 80 | 173 |
| 66 | D:F | TCDMDA:BDDA | 79 | 45 |
| 67 | B:D | NGDA:TCDMDA | 78 | 97 |
| 68 | H:P | GDMA:THFuA | 78 | 72 |
| 69 | M:Q | EGDPEA:BA | 78 | 98 |
| 70 | M:I | EGDPEA:DEAEMA | 77 | 89 |
| 71 | A:G | HBOPBA:EGDA | 76 | 74 |
| 72 | C:D | BHMOPhP:TCDMDA | 76 | 83 |
| 73 | H:A | GDMA:HBOPBA | 73 | 64 |
| 74 | B:O | NGDA:MAETA | 72 | 45 |
| 75 | N:H | FuMA:GDMA | 72 | 48 |
| 76 | N:E | FuMA:EG4DMA | 72 | 47 |
| 77 | F:M | BDDA:EGDPEA | 70 | 82 |
| 78 | F | BDDA | 70 | 68 |
| 79 | M:R | EGDPEA:TDFOMA | 69 | 116 |
| 80 | O:F | MAETA:BDDA | 68 | 57 |
| 81 | B:I | NGDA:DEAEMA | 68 | 40 |
| 82 | P:Q | THFuA:BA | 68 | 107 |
| 83 | I:D | DEAMEA:TCDMDA | 68 | 38 |
| 84 | G:O | EGDA:MAETA | 67 | 75 |
| 85 | B:M | NGDA:EGDPEA | 67 | 75 |
| 86 | H:E | GDMA:EG4DMA | 66 | 54 |
| 87 | L:M | HEMA:EGDPEA | 65 | 68 |
| 88 | A:C | HBOPBA:BHMOPhP | 65 | 63 |
| 89 | F:D | BDDA:TCMDMA | 65 | 64 |
| 90 | F:I | BDDA:DEAEMA | 65 | 45 |
| 91 | H:O | GDMA:MAETA | 64 | 80 |
| 92 | O:E | MAETA:EG4DMA | 63 | 55 |
| 93 | M:F | EGDPEA:BDDA | 62 | 69 |
| 94 | J:B | mMAOES:NGDA | 62 | 110 |
| 95 | D:L | TCMDMA:HEMA | 62 | 51 |
| 96 | B | NGDA | 62 | 75 |
| 97 | F:B | BDDA:NGDA | 62 | 51 |
| 98 | H:I | GDMA:DEAEMA | 61 | 76 |
| 99 | C:B | BHMOPhP:NGDA | 61 | 74 |
| 100 | D:M | TCMDMA:EGDPEA | 61 | 74 |
| 101 | F:L | BDDA:HEMA | 60 | 58 |
| 102 | D:A | TCMDMA:HBOPBA | 60 | 48 |
| 103 | E:F | EG4DMA:BDDA | 59 | 63 |
| 104 | G:A | EGDA:HBOPBA | 59 | 129 |
| 105 | B:G | NGDA:EGDA | 59 | 71 |
| 106 | N:Q | FuMA:BA | 59 | 127 |
| 107 | H:M | GDMA:EGDPEA | 59 | 45 |
| 108 | O:G | MAETA:EGDA | 59 | 64 |
| 109 | E:B | EG4DMA:NGDA | 58 | 64 |
| 110 | M:H | EGDPEA:GDMA | 57 | 76 |
| 111 | E:G | EG4DMA:EGDA | 57 | 56 |
| 112 | A | HBOPBA | 56 | 43 |
| 113 | A:F | HBOPBA:BDDA | 56 | 58 |
| 114 | D:H | TCMDMA:GDMA | 56 | 52 |
| 115 | N:K | FuMA:tBAEMA | 56 | 38 |
| 116 | F:C | BDDA:BHMOPhP | 56 | 56 |
| 117 | A:D | HBOPBA:TCMDMA | 56 | 53 |
| 118 | F:R | BDDA:TDFOMA | 55 | 46 |
| 119 | G:J | EGDA:mMAOES | 55 | 67 |
| 120 | D:I | TCMDMA:DEAEMA | 55 | 49 |
| 121 | P:G | THFuA:EGDA | 55 | 73 |
| 122 | L:A | HEMA:HBOPBA | 55 | 56 |
| 123 | A:E | HBOPBA:EG4DMA | 55 | 70 |
| 124 | M:D | EGDPEA:TCMDMA | 54 | 53 |
| 125 | M:K | EGDPEA:tBAEMA | 54 | 71 |
| 126 | Q:D | BA:TCMDMA | 54 | 63 |
| 127 | I:F | DEAMEA:BDDA | 54 | 70 |
| 128 | J:D | mMAOES:TCMDMA | 53 | 37 |
| 129 | K:O | tBAEMA:MAETA | 53 | 41 |
| 130 | R:F | TDFOMA:BDDA | 53 | 65 |
| 131 | M:A | EGDPEA:HBOPBA | 53 | 45 |
| 132 | A:H | HBOPBA:GDMA | 52 | 84 |
| 133 | Q:B | BA:NGDA | 52 | 52 |
| 134 | B:C | NGDA:BHMOPhP | 52 | 25 |
| 135 | E:C | EG4DMA:BHMOPhP | 52 | 51 |
| 136 | I:H | DEAMEA:GDMA | 51 | 122 |
| 137 | O:M | MAETA:EGDPEA | 51 | 62 |
| 138 | C:J | BHMOPhP:mMAOES | 49 | 52 |
| 139 | C:L | BHMOPhP:HEMA | 48 | 84 |
| 140 | C:F | BHMOPhP:BDDA | 48 | 61 |
| 141 | M:O | EGDPEA:MAETA | 48 | 89 |
| 142 | C:E | BHMOPhP:EG4DMA | 47 | 49 |
| 143 | D:E | TCDMDA:EG4DMA | 47 | 50 |
| 144 | E:J | EG4DMA:mMAOES | 47 | 56 |
| 145 | K:M | tBAEMA:EGDPEA | 47 | 74 |
| 146 | R:A | TDFOMA:HBOPBA | 46 | 77 |
| 147 | G:R | EGDA:TDFOMA | 45 | 106 |
| 148 | C:O | BHMOPhP:MAETA | 45 | 57 |
| 149 | Q:G | BA:EGDA | 45 | 53 |
| 150 | P:M | THFuA:EGDPEA | 45 | 42 |
| 151 | I:G | DEAMEA:EGDA | 44 | 37 |
| 152 | I:Q | DEAMEA:BA | 44 | 62 |
| 153 | M:L | EGDPEA:HEMA | 44 | 41 |
| 154 | C:G | BHMOPhP:EGDA | 44 | 40 |
| 155 | G:E | EGDA:EG4DMA | 43 | 51 |
| 156 | E:D | EG4DMA:TCDMDA | 43 | 47 |
| 157 | B:S | NGDA:HMAm | 42 | 71 |
| 158 | I:N | DEAMEA:FuMA | 42 | 44 |
| 159 | O:N | MAETA:FuMA | 41 | 51 |
| 160 | N:A | FuMA:HBOPBA | 41 | 37 |
| 161 | O:Q | MAETA:BA | 41 | 41 |
| 162 | H:L | GDMA:HEMA | 40 | 30 |
| 163 | E | EG4DMA | 40 | 49 |
| 164 | H | GDMA | 40 | 38 |
| 165 | E:S | EG4DMA:HMAm | 40 | 59 |
| 166 | F:S | BDDA:HMAm | 40 | 51 |
| 167 | G:B | EGDA:NGDA | 38 | 41 |
| 168 | Q:C | BA:BHMOPhP | 38 | 47 |
| 169 | L:G | HEMA:EGDA | 37 | 50 |
| 170 | P:A | THFuA:HBOPBA | 36 | 31 |
| 171 | B:E | NGDA:EG4DMA | 36 | 25 |
| 172 | O:P | MAETA:THFuA | 36 | 32 |
| 173 | N:B | FuMA:NGDA | 36 | 40 |
| 174 | A:J | HBOPBA:mMAOES | 35 | 45 |
| 175 | P:S | THFuA:HMAm | 35 | 28 |
| 176 | L:O | HEMA:MAETA | 34 | 47 |
| 177 | Q:A | BA:HBOPBA | 34 | 48 |
| 178 | R:H | TDFOMA:GDMA | 34 | 54 |
| 179 | M:N | EGDPEA:FuMA | 34 | 76 |
| 180 | D:S | TCDMDA:HMAm | 34 | 35 |
| 181 | K:A | tBAEMA:HBOPBA | 34 | 79 |
| 182 | H:J | GDMA:mMAOES | 33 | 58 |
| 183 | C:S | BHMOPhP:HMAm | 33 | 67 |
| 184 | E:H | EG4DMA:GDMA | 33 | 53 |
| 185 | R:D | TDFOMA:TCDMDA | 33 | 26 |
| 186 | L:I | HEMA:DEAEMA | 32 | 39 |
| 187 | N:P | FuMA:THFuA | 32 | 31 |
| 188 | K:B | tBAEMA:NGDA | 32 | 39 |
| 189 | K:E | tBAEMA:EG4DMA | 31 | 46 |
| 190 | P:O | THFuA:MAETA | 31 | 40 |
| 191 | O | MAETA | 31 | 15 |
| 192 | Q:O | BA:MAETA | 31 | 25 |
| 193 | B:A | NGDA:HBOPBA | 31 | 50 |
| 194 | B:P | NGDA:THFuA | 30 | 34 |
| 195 | K:L | tBAEMA:HEMA | 30 | 23 |
| 196 | R:N | TDFOMA:FuMA | 29 | 21 |
| 197 | K:P | tBAEMA:THFuA | 29 | 24 |
| 198 | Q | BA | 29 | 59 |
| 199 | E:O | EG4DMA:MAETA | 29 | 39 |
| 200 | O:S | MAETA:HMAm | 29 | 20 |
| 201 | C:I | BHMOPhP:DEAEMA | 28 | 57 |
| 202 | K:F | tBAEMA:BDDA | 28 | 38 |
| 203 | Q:R | BA:TDFOMA | 28 | 37 |
| 204 | S:F | HMAm:BDDA | 28 | 67 |
| 205 | Q:H | BA:GDMA | 27 | 48 |
| 206 | N:G | FuMA:EGDA | 27 | 32 |
| 207 | Q:F | BA:BDDA | 27 | 27 |
| 208 | A:I | HBOPBA:DEAEMA | 26 | 40 |
| 209 | N:L | FuMA:HEMA | 26 | 34 |
| 210 | K:Q | tBAEMA:BA | 26 | 23 |
| 211 | J:E | mMAOES:EG4DMA | 26 | 23 |
| 212 | E:Q | EG4DMA:BA | 26 | 46 |
| 213 | R:E | TDFOMA:EG4DMA | 25 | 24 |
| 214 | A:S | HBOPBA:HMAm | 25 | 30 |
| 215 | J:F | mMAOES:BDDA | 25 | 34 |
| 216 | C:Q | BHMOPhP:BA | 24 | 40 |
| 217 | K:C | tBAEMA:BHMOPhP | 23 | 25 |
| 218 | J:G | mMAOES:EGDA | 23 | 31 |
| 219 | A:N | HBOPBA:FuMA | 23 | 52 |
| 220 | P:J | THFuA:mMAOES | 22 | 25 |
| 221 | Q:N | BA:FuMA | 22 | 25 |
| 222 | I:C | DEAMEA:BHMOPhP | 22 | 27 |
| 223 | I:A | DEAMEA:HBOPBA | 21 | 20 |
| 224 | Q:M | BA:EGDPEA | 21 | 40 |
| 225 | A:M | HBOPBA:EGDPEA | 21 | 36 |
| 226 | E:K | EG4DMA:tBAEMA | 21 | 41 |
| 227 | R:P | TDFOMA:THFuA | 21 | 22 |
| 228 | I:K | DEAMEA:tBAEMA | 21 | 21 |
| 229 | I:M | DEAMEA:EGDPEA | 21 | 20 |
| 230 | C:N | BHMOPhP:FuMA | 20 | 25 |
| 231 | H:S | GDMA:HMAm | 20 | 26 |
| 232 | Q:P | BA:THFuA | 20 | 35 |
| 233 | E:M | EG4DMA:EGDPEA | 20 | 52 |
| 234 | L:S | HEMA:HMAm | 19 | 27 |
| 235 | S:G | HMAm:EGDA | 19 | 16 |
| 236 | G:M | EGDA:EGDPEA | 19 | 29 |
| 237 | D:K | TCDMDA:tBAEMA | 19 | 24 |
| 238 | C:H | BHMOPhP:GDMA | 19 | 21 |
| 239 | S:N | HMAm:FuMA | 18 | 17 |
| 240 | F:A | BDDA:HBOPBA | 18 | 16 |
| 241 | Q:L | BA:HEMA | 18 | 20 |
| 242 | P:L | THFuA:HEMA | 18 | 20 |
| 243 | G:S | EGDA:HMAm | 18 | 23 |
| 244 | M:P | EGDPEA:THFuA | 18 | 34 |
| 245 | J | mMAOES | 18 | 34 |
| 246 | P:I | THFuA:DEAEMA | 18 | 11 |
| 247 | S:E | HMAm:EG4DMA | 18 | 21 |
| 248 | J:A | mMAOES:HBOPBA | 17 | 23 |
| 249 | I:O | DEAMEA:MAETA | 17 | 16 |
| 250 | K:G | tBAEMA:EGDA | 17 | 19 |
| 251 | L:F | HEMA:BDDA | 17 | 24 |
| 252 | K:R | tBAEMA:TDFOMA | 17 | 13 |
| 253 | I:L | DEAMEA:HEMA | 17 | 14 |
| 254 | A:P | HBOPBA:THFuA | 17 | 34 |
| 255 | L:P | HEMA:THFuA | 17 | 21 |
| 256 | B:Q | NGDA:BA | 16 | 21 |
| 257 | E:I | EG4DMA:DEAEMA | 16 | 15 |
| 258 | R:M | TDFOMA:EGDPEA | 16 | 27 |
| 259 | K:N | tBAEMA:FuMA | 16 | 13 |
| 260 | I:R | DEAMEA:TDFOMA | 16 | 36 |
| 261 | R:J | TDFOMA:mMAOES | 16 | 19 |
| 262 | B:K | NGDA:tBAEMA | 16 | 18 |
| 263 | M:S | EGDPEA:HMAm | 16 | 23 |
| 264 | C:A | BHMOPhP:HBOPBA | 16 | 28 |
| 265 | M:G | EGDPEA:EGDA | 16 | 26 |
| 266 | R:S | TDFOMA:HMAm | 16 | 19 |
| 267 | E:N | EG4DMA:FuMA | 16 | 25 |
| 268 | S:D | HMAm:TCDMDA | 16 | 15 |
| 269 | L:C | HEMA:BHMOPhP | 16 | 20 |
| 270 | A:O | HBOPBA:MAETA | 15 | 28 |
| 271 | L:H | HEMA:GDMA | 15 | 26 |
| 272 | M | EGDPEA | 15 | 18 |
| 273 | K | tBAEMA | 15 | 8 |
| 274 | S:M | HMAm:EGDPEA | 15 | 7 |
| 275 | J:P | mMAOES:THFuA | 15 | 18 |
| 276 | K:H | tBAEMA:GDMA | 14 | 14 |
| 277 | G:I | EGDA:DEAEMA | 14 | 19 |
| 278 | R:K | TDFOMA:tBAEMA | 14 | 12 |
| 279 | G:H | EGDA:GDMA | 14 | 15 |
| 280 | A:Q | HBOPBA:BA | 13 | 23 |
| 281 | K:I | tBAEMA:DEAEMA | 13 | 10 |
| 282 | S:K | HMAm:tBAEMA | 13 | 27 |
| 283 | P:K | THFuA:tBAEMA | 13 | 23 |
| 284 | J:M | mMAOES:EGDPEA | 13 | 11 |
| 285 | B:N | NGDA:FuMA | 13 | 11 |
| 286 | O:R | MAETA:TDFOMA | 13 | 13 |
| 287 | R:O | TDFOMA:MAETA | 12 | 9 |
| 288 | G:K | EGDA:tBAEMA | 12 | 19 |
| 289 | R:L | TDFOMA:HEMA | 12 | 10 |
| 290 | R:Q | TDFOMA:BA | 11 | 13 |
| 291 | S:C | HMAm:BHMOPhP | 11 | 11 |
| 292 | L:N | HEMA:FuMA | 11 | 8 |
| 293 | R | TDFOMA | 11 | 11 |
| 294 | I | DEAEMA | 11 | 12 |
| 295 | O:L | MAETA:HEMA | 11 | 13 |
| 296 | G:L | EGDA:HEMA | 10 | 12 |
| 297 | S:O | HMAm:MAETA | 10 | 11 |
| 298 | C:P | BHMOPhP:THFuA | 10 | 9 |
| 299 | L:Q | HEMA:BA | 10 | 9 |
| 300 | N:R | FuMA:TDFOMA | 10 | 14 |
| 301 | N:M | FuMA:EGDPEA | 10 | 13 |
| 302 | N:S | FuMA:HMAm | 10 | 10 |
| 303 | K:D | tBAEMA:TCDMDA | 10 | 9 |
| 304 | S:J | HMAm:mMAOES | 9 | 7 |
| 305 | J:N | mMAOES:FuMA | 9 | 16 |
| 306 | J:C | mMAOES:BHMOPhP | 9 | 9 |
| 307 | I:J | DEAMEA:mMAOES | 9 | 10 |
| 308 | S:L | HMAm:HEMA | 9 | 7 |
| 309 | O:I | MAETA:DEAEMA | 9 | 16 |
| 310 | S:B | HMAm:NGDA | 9 | 8 |
| 311 | S:I | HMAm:DEAEMA | 9 | 5 |
| 312 | I:S | DEAEMA:HMAm | 9 | 8 |
| 313 | M:J | EGDPEA:mMAOES | 9 | 8 |
| 314 | C:M | BHMOPhP:EGDPEA | 8 | 12 |
| 315 | S:H | HMAm:GDMA | 8 | 6 |
| 316 | A:K | HBOPBA:tBAEMA | 8 | 13 |
| 317 | G:N | EGDA:FuMA | 8 | 7 |
| 318 | S:P | HMAm:THFuA | 8 | 14 |
| 319 | L:K | HEMA:tBAEMA | 8 | 6 |
| 320 | K:S | tBAEMA:HMAm | 8 | 6 |
| 321 | E:L | EG4DMA:HEMA | 8 | 16 |
| 322 | R:C | TDFOMA:BHMOPhP | 8 | 10 |
| 323 | J:K | mMAOES:tBAEMA | 8 | 7 |
| 324 | Q:K | BA:tBAEMA | 8 | 6 |
| 325 | S:A | HMAm:HBOPBA | 8 | 9 |
| 326 | Q:J | BA:mMAOES | 7 | 13 |
| 327 | L | HEMA | 7 | 6 |
| 328 | Q:S | BA:HMAm | 7 | 12 |
| 329 | O:K | MAETA:tBAEMA | 7 | 10 |
| 330 | R:I | TDFOMA:DEAEMA | 7 | 5 |
| 331 | A:L | HBOPBA:HEMA | 7 | 7 |
| 332 | O:J | MAETA:mMAOES | 6 | 9 |
| 333 | L:J | HEMA:mMAOES | 6 | 7 |
| 334 | E:R | EG4DMA:TDFOMA | 6 | 15 |
| 335 | F:Q | BDDA:BA | 6 | 8 |
| 336 | E:P | EG4DMA:THFuA | 6 | 9 |
| 337 | K:J | tBAEMA:mMAOES | 6 | 8 |
| 338 | F:N | BDDA:FuMA | 6 | 7 |
| 339 | N:O | FuMA:MAETA | 6 | 8 |
| 340 | G:P | EGDA:THFuA | 6 | 5 |
| 341 | Q:I | BA:DEAEMA | 6 | 5 |
| 342 | O:C | MAETA:BHMOPhP | 5 | 7 |
| 343 | J:H | mMAOES:GDMA | 5 | 8 |
| 344 | J:O | mMAOES:MAETA | 5 | 8 |
| 345 | L:R | HEMA:TDFOMA | 5 | 6 |
| 346 | S:R | HMAm:TDFOMA | 4 | 4 |
| 347 | N:J | FuMA:mMAOES | 4 | 6 |
| 348 | S:Q | HMAm:BA | 4 | 4 |
| 349 | G:Q | EGDA:BA | 4 | 5 |
| 350 | J:L | mMAOES:HEMA | 3 | 4 |
| 351 | J:S | mMAOES:HMAm | 3 | 6 |
| 352 | I:P | DEAMEA:THFuA | 3 | 3 |
| 353 | J:I | mMAOES:DEAEMA | 3 | 4 |
| 354 | D:R | TCDMDA:TDFOMA | 3 | 4 |
| 355 | J:Q | mMAOES:BA | 2 | 3 |
| 356 | J:R | mMAOES:TDFOMA | 2 | 3 |
| 357 | S | HMAM | 2 | 3 |
| 358 | C:K | BHMOPhP:tBAEMA | 2 | 1 |
| 359 | C:R | BHMOPhP:TDFOMA | 2 | 3 |

**Table S4:**
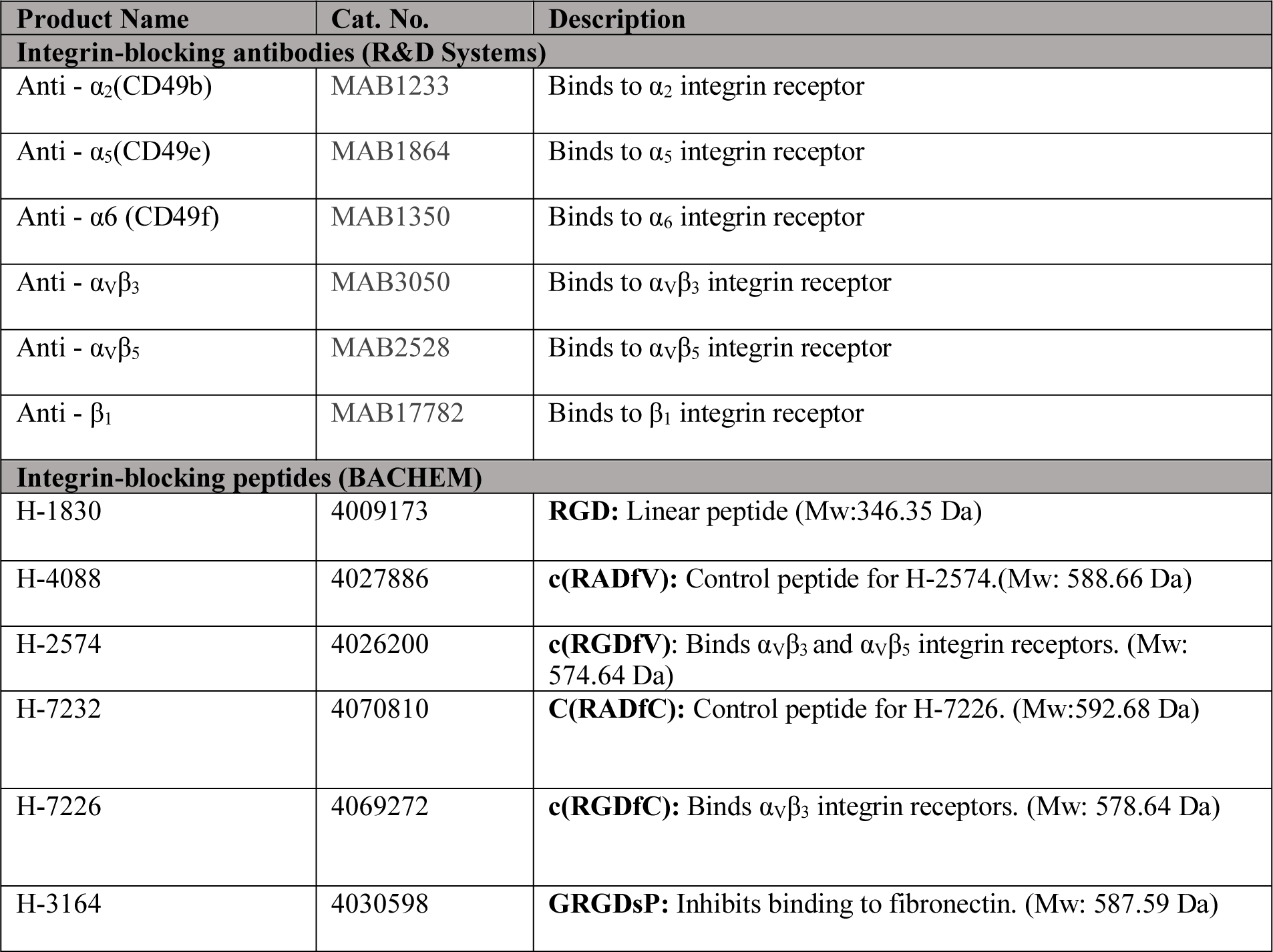
Integrin blocking antibodies and peptides.

## Notes

### Competing Interest Statement

The authors have declared no competing interest.

